# Genome-wide association, prediction and heritability in bacteria

**DOI:** 10.1101/2021.10.04.462983

**Authors:** Sudaraka Mallawaarachchi, Gerry Tonkin-Hill, Nicholas J. Croucher, Paul Turner, Doug Speed, Jukka Corander, David Balding

## Abstract

Advances in whole-genome genotyping and sequencing have allowed genome-wide analyses of association, prediction and heritability in many organisms. However, the application of such analyses to bacteria is still in its infancy, being limited by difficulties including the plasticity of bacterial genomes and their strong population structure. Here we propose, and validate using simulations, a suite of genome-wide analyses for bacteria. We combine methods from human genetics and previous bacterial studies, including linear mixed models, elastic net and LD-score regression, and introduce innovations such as frequency-based allele coding, testing for both insertion/deletion and nucleotide effects and partitioning heritability by genome region. We then analyse three phenotypes of a major human pathogen *Streptococcus pneumoniae*, including the first analyses of minimum inhibitory concentrations (MIC) for each of two antibiotics, penicillin and ceftriaxone. We show that these are highly heritable leading to high prediction accuracy, which is explained by many genetic associations identified under good control of population structure effects. In the case of ceftriaxone MIC, these results are surprising because none of the isolates was resistant according to the inhibition zone diameter threshold. We estimate that just over half of the heritability of penicillin MIC is explained by a known drug-resistance region, which also contributes around a quarter of the heritability of ceftriaxone MIC. For the within-host survival phenotype carriage duration, no reliable associations were found but we observed moderate heritability and prediction accuracy, indicating a polygenic trait. While generating important new results for *S. pneumoniae*, we have critically assessed existing methods and introduced innovations that will be useful for future large-scale population genomics studies to help decipher the genetic architecture of bacterial traits.

**Author summary:** Genome-wide association, prediction and heritability analyses in bacteria are beginning to help unravel the genetic underpinnings of traits such as antimicrobial resistance, virulence, within-host survival and transmissibility. Progress to date is limited by challenges including the effects of strong population structure and variable recombination, and the many gaps in sequence alignments including the absence of entire genes in many isolates. More work is required to critically asses and develop methods for bacterial genomics. We address this task here, using a range of existing methods from bacterial and human genetics, such as linear mixed models, elastic net and LD-score regression. Using simulations, we first validate and then adapt these methods to introduce new analyses, including separate assessment of gap and nucleotide effects, a new allele coding for association analyses and a method to partition heritability into genome regions. We analyse within-host survival and two antimicrobial response traits of *Streptococcus pneumoniae*, identifying many novel associations while demonstrating good control of population structure and accurate prediction. We present both new results for an important pathogen and methodological advances that will be useful in guiding future studies in bacterial population genomics.

## Introduction

The ability to perform genome-wide analyses of DNA variations has enabled detailed investigations of the genetic architecture of traits in many organisms. In human genetics, the study of heritability across the genome has received considerable attention and the main statistical challenges related to robust estimation of SNP heritability are being overcome [1, 2]. Similar studies in bacteria are emerging [3, 4], but the pros and cons of the many available methods have not yet been extensively studied. We adopted popular methods from human genetics, using linear mixed models (LMMs) and linkage disequilibrium score regression (LDSC) to investigate genome-wide association and heritability, in combination with elastic-net regression for prediction of three traits (two not previously studied) in *Streptococcus pneumoniae*.

*S. pneumoniae*, or the pneumococcus, is a Gram-positive human pathogen that can cause several invasive diseases such as pneumonia, meningitis and sepsis, as well as milder diseases such as acute otitis media and tonsillitis. Typically, pneumococci colonise the nasopharynx of a host asymptomatically and transmit effectively between young children, who frequently carry the bacterium until they develop broad natural immunity. This may be supplemented by vaccination with any of the polysaccharide conjugate vaccines (PCVs), which induce effective protection against some common virulent serotypes. Several population genomic studies have characterized central epidemiological traits of the pneumococcus, including duration of carriage and resistance to commonly used antibiotics.

In a pioneering study, Lees et al. [3], found high heritability of the duration of carriage of *S. pneumoniae* in human hosts. Furthermore, the strong genetic control of the binary trait antimicrobial resistance (AMR) is well established from genome-wide association studies (GWAS) [5–8]. The quantitative trait minimum inhibitory concentration (MIC) has previously been studied in *Mycobacterium tuberculosis* [9], but not in *S. pneumoniae*.

We critically assess available methods for association, prediction and heritability analyses, and propose novel developments, which we use to investigate carriage duration (CD), ceftriaxone MIC and penicillin MIC in *S. pneumoniae*, finding many new associations and high predictive accuracy for the two MIC traits. Given the increasing availability of large-scale bacterial GWAS, the developments presented here will provide a useful guide to future studies.

## Materials and methods

### Source of data

The present study is based on nasopharyngeal swab data collected monthly from infants and their mothers in the Maela refugee camp in Thailand between 2007 and 2010 [10]. Overall, 23 910 swabs were collected during the original cohort study, from which 19 359 swabs from 737 infants and 952 mothers were processed according to World Health Organization (WHO) pneumococcal carriage detection protocols [11] and/or the latex sweep method [12].

Penicillin and ceftriaxone susceptibilities were assessed using 1 μg oxacillin disks in accordance with the 2007 CLSI guidelines [13]. Only isolates with an oxacillin zone diameter of <20 mm were subject to benzyl penicillin and ceftriaxone MIC measurements; other isolates were classified as susceptible.

### Preparation of phenotypes

Following [3], we implemented a hidden Markov model, using the R package msm [14], to obtain maximum-likelihood estimates of CD values. Due to differences in immune response to bacterial infections between adults and infants [15], only data from infants were used for CD analyses, but we analysed all MIC values regardless of the host. To obtain approximate normal distributions, we log-transformed all three phenotypes (see S1 Fig for histograms).

### Preparation of genetic data

We used a published dataset [5] of high quality genome sequences from 2 663 isolates, manually selected and aligned to the ATCC700669 reference genome using the snippy pipeline version 4.4.0 [16], with minimum coverage set at the default 10 reads. Of these, 1 612 isolates were sampled during *S. pneumoniae* positive episodes, on average 1.5 (SD 1.0) isolates per episode. For the 337 episodes represented by *>* 1 genome sequence, we used the sequence from the last isolate sampled. This resulted in 1 047 sequenced CD episodes in 370 host infants (mean 2.8, SD 1.9 episodes per host). The median CD was 64 days, with mean 110 and SD 102. MIC data for both penicillin (mean 0.57, SD 0.48 μg ml^−1^) and ceftriaxone (mean 0.36, SD 0.28 μg ml^*−*1^) were available for 1 332 isolates, of which 554 also have a CD episode. SNP-sites version 2.5.1 [17] and VCFtools version 0.1.16 [18] were used to identify 239 176 variant sites in the CD dataset, and 215 892 in the MIC dataset.

A gene was considered a part of the core genome if it was observed in ≥ 95% of isolates, otherwise it was labelled as *accessory*. Pangenome data were extracted by assembling and annotating the read sequences using Prokka version 1.14.6 [19]. Orthologous and paralogous gene clusters were then inferred using the Panaroo pangenome pipeline version 1.2.4, generating a gene presence/absence matrix [20]. While the core genome was analysed at each variant site, the accessory genome was analysed at the level of genes, using standardised gene counts. The numbers of accessory genes showing variation in the CD and MIC datasets, respectively, were 2 310 and 2 242.

### Association analyses

#### Testing gap and SNP effects

Five alleles are possible at each variant site, the four nucleotides and gap. Gaps are observed at approximately 71% of variant sites (see Fig. 1 for the gap frequency distribution), while two, three and four nucleotide alleles are observed at 71%, 7% and 0.4% of variant sites, respectively. In human genetics, multi-allelic SNPs and gaps are both rare and SNP alleles are usually coded as binary, leading to three diploid genotypes that can be coded using two degrees of freedom (df), or 1 df under an additive model. For haploid bacteria, a general coding would require up to 4 df per SNP. The usual approach in previous analyses is a 1 df binary coding indicating presence/absence of the major allele. This coding loses information if the minor alleles have different effects. In particular, gap and SNP effects can differ, due in part to different local-dependence effects of insertion/deletion lengths and recombination.

**Fig 1.**
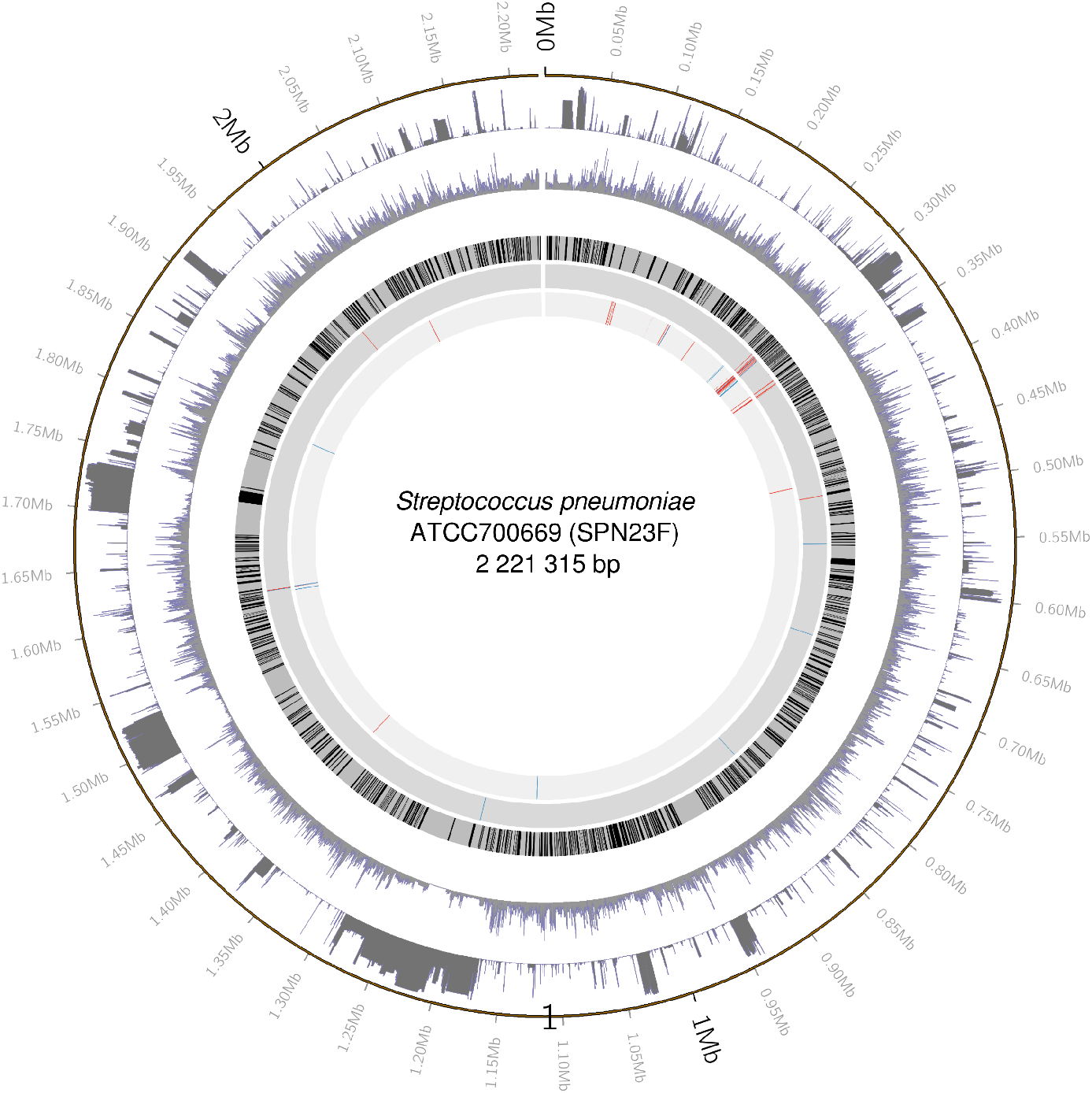
Mapping of association hits to the ATCC700669 reference genome. Working inwards from the outer circle showing basepair positions along the genome, the subsequent circles show the distributions of gap and minor allele frequencies in the MIC dataset, annotated core genes (in black), and SNPs associated with ceftriaxone MIC and penicillin MIC according to the gap test (blue) and SNP test (red). Figure prepared using circos [21].

In previous bacterial GWAS analyses, variant sites with many gaps have often been removed. Reasons include that a gap coding can reflect data quality issues other than a true insertion/deletion sequence state, and that the effects of large insertions or deletions cannot be localised to specific sites. However, insertions and deletions that generate gaps can affect phenotypes, and it is of interest to identify them, while recognising that the ultimate cause of the association signal may be difficult to decipher. For the core genome variants, we first used a binary gap/non-gap coding to compute a ‘gap test’ statistic at sites with ≥ 10 of both gap and non-gap sequences. The test statistic at the *j*th variant was the squared standardised effect size: 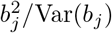. Next we computed a ‘SNP test’ statistic, omitting gap sequences, at sites with ≥ 10 copies of at least two nucleotides. We used a 1 df allele coding equal to the sample frequency of the allele, which assumes that effect sizes vary linearly with allele frequency. For sites with both gap and SNP statistics available, the larger one was used (“max” statistic). In the simulation study we also combined the two statistics using Stouffer’s method (divide the sum by 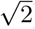), which we refer to as the “combi” statistic.

To ensure a family-wise error rate (FWER) of 0.05, we performed 500 permutations of the ceftriaxone MIC phenotype, each time re-running the association analysis pipeline and recording the maximum test statistic. Our significance threshold for the real-data analyses was 24.8, the 25th largest of the 500 maximum test statistics. In comparison, the corresponding Bonferroni threshold based on 133K tests and a 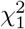 null distribution, is 25.8. Therefore, while taking the max of gap and SNP test statistics tends to inflate the null distribution, Bonferroni correction would still be conservative because it ignores the correlations among the statistics. Because of the similarity of the phenotype distributions (S1 Fig), for penicillin MIC we used the permutation threshold derived for ceftriaxone MIC.

For comparison, we also employed a 1 df association test based on presence/absence of the major allele at each variant, whether gap or a nucleotide, using the Bonferroni threshold. While this test allows some gap effects to be detected, if gap is not the major allele it assumes that the gap and minor nucleotide effects are the same. If gap is the major allele then all nucleotide effects are assumed to be the same.

#### Population structure, phylogeny and clustering

Levels of recombination vary over bacterial species, but in general asexual reproduction leads to strong population structure, which is challenging for association analyses [22, 23]. Population structure refers to groups of individuals (sub-populations) with greater genetic similarity among them than with other individuals, which causes genome-wide genetic correlations that can confound association signals. Sub-populations may also differ in environmental exposures, which can compound the problem.

There is no complete solution to the problems caused by population structure, and attempts to address them risk discarding true as well as spurious signal. Most approaches introduce either covariates or a genetic random effect into association models to absorb signals that can be explained by population structure, which then do not contribute to association statistics. The variance-covariance matrix **G** of a genetic random effect is assumed known *a priori* based on measures of similarity between pairs of sequences.

Sequence clusters can be used to define either **G**, via cluster distances, or population structure covariates via indicators of cluster membership. Clustering can proceed by constructing a phylogenetic tree that models the evolutionary history of the sequences [24], with nodes of the tree used as cluster identifiers and branch lengths used to define cluster distances. We inferred maximum-likelihood phylogenies of both CD and MIC datasets using IQTree version 2.0.6 [25] under the general time reversible model, with discrete Gamma (+G option) base substitution rates across sites (Fig. 2). The model assumes no recombination, which is false for *S. pneumoniae*, and consequently the usefulness of the resulting phylogeny has been questioned [26].

**Fig 2.**
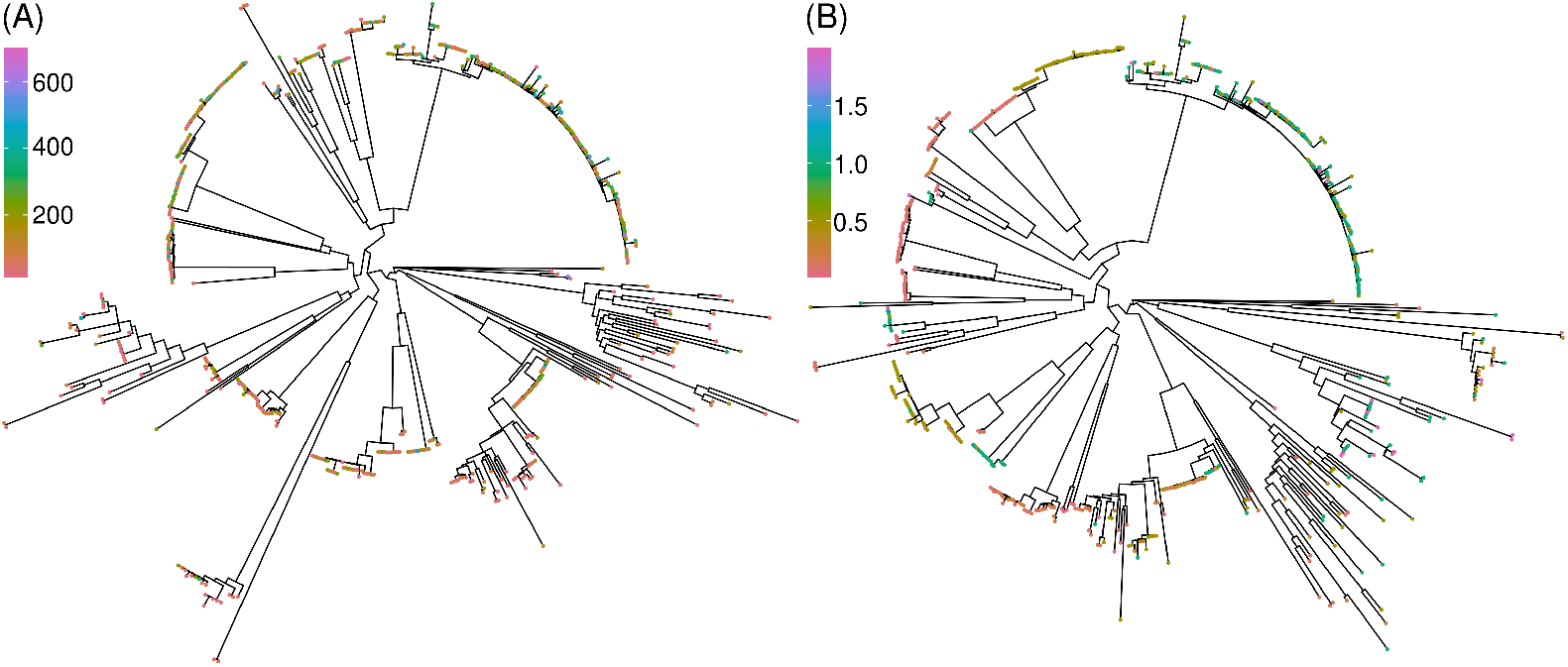
Phylogenies inferred using IQtree2. (A) 1 047 isolates with a carriage duration (CD) phenotype, indicated by tip colour (in days). (B) 1 332 isolates with MIC phenotypes, with the penicillin phenotype indicated by tip colour (in μg ml^*−*1^). Plots generated after midpoint rooting using R packages ape [27], phytools [28] and ggtree [29].

FastBAPS, which extends hierBAPS, [30–32] was also used to cluster the isolates, without reference to a phylogeny. This approach generates an initial clustering using between-variant pairwise distances based on Ward’s method [33], then an optimal set of clusters is identified using Bayesian hierarchical clustering [34].

In human studies, **G** was in the past computed from known pedigrees [35] and now usually as a genome-wide average allelic correlation [36]. For bacteria, **G** can be defined using allelic correlations under any 1 df allele coding. Despite the success of this approach in human studies, our preliminary analyses could not identify an allele coding that led to good control of population structure effects, although using the gap presence/absence binary indicator gave the best results among those we tried.

Conversely, despite the questionable validity of the phylogeny due to it ignoring recombination, defining **G** in terms of lengths of shared phylogenetic branches [37] led to good control of population structure, as evidenced by QQ plots.

#### Linear mixed model (LMM) analyses

We wish to test *b*_*j*_ = 0 within the LMM [38]:

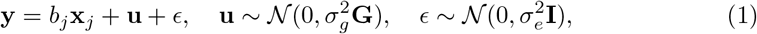

where **y** is a length-*n* phenotype vector, **x**_*j*_ is the vector encoding alleles at the *j*th variant, and **u** and *ϵ* are random vectors of genetic and environmental effects, with **I** the *n* × *n* identity matrix.

Pyseer [39] has recently been widely used in bacterial GWAS, and an extensive summary of its models with performance benchmarking is available [40]. The Pyseer implementation of (1) is based on FaST-LMM [41], and includes likelihood ratio testing of *b*_*j*_ = 0. It requires binary coding of genetic variants, and so can be used for the gap and major-allele tests, but it cannot accommodate the frequency-coding or omission of the gap sequences at each SNP test. To overcome this problem, we used a two-stage LMM/GLS pipeline for the SNP test, similar to EMMAX [42], in which the phenotype for association testing was the residual from fitting (1) with *b*_*j*_ = 0. This first ‘LMM’ stage was performed using lme4qtl [35]. The *b*_*j*_ were then estimated in a second stage using generalised least squares regression (GLS). In the CD analyses for the SNP test, we were able to incorporate an extra random effect to model shared host in the LMM/GLS pipeline, but for the gap and MA tests performed using Pyseer-LMM, this was replaced by a binary covariate indicating previous carriage.

Accessory genome genes were tested using the LMM/GLS pipeline, with a single test based on standardised gene counts.

#### Phylogenetic method treeWAS

For comparison, we also implemented the phylogeny-based treeWAS [43] using the MA coding. Use of a single phylogeny in treeWAS corresponds to an assumption of negligible recombination. As recommended for recombinant species such as *S. pneumoniae* [43], we first implemented the ClonalFrameML pipeline (see S2 Fig) [44]. Then treeWAS infers the ancestral phenotype and genotype states at each internal node of the phylogeny, before computing three association test statistics:

1. **Terminal Score**: measures sample-wide phenotype-genotype associations between leaves of the phylogeny.
2. **Simultaneous Score**: measures parallel changes in both phenotype and genotype on phylogeny branches.
3. **Subsequent Score**: measures the proportion of the tree within which genotype and phenotype ‘co-exist’. It is equivalent to integrating association scores over all tree nodes.

For each sore, a significance threshold was estimated from null simulations of genetic data at 10 times as many sites as the observed dataset.

### Phenotype prediction: whole genome elastic net (wg-enet)

We set up the Pyseer wg-enet model in glmnet [45] in order to use a frequency-based allele coding as in the SNP test except that gaps were counted as an allele. Following Pyseer guidelines [46], we omitted 25% of variants with the largest association *p*-values, and then removed highly-correlated variants at a 0.75 threshold. We verified the finding of [46] that prediction accuracy is improved using weight *w*_*i*_ for the *i*th isolate, where *w*_*i*_ is proportional to the inverse of the size of the cluster that includes the isolate, and Σ_*i*_ *w*_*i*_ = *n*. After centering the phenotype values to have mean zero, the *i*th phenotype value is predicted by 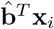, where **x**_*i*_ is the vector of allele indicators for the *i*th sequence, and

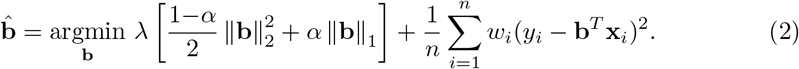

We use cross-validation (CV) to optimise the penalty parameter *λ*. When *λ* = 0 we have weighted least-squares regression, while increasing *λ* reduces overfitting but introduces bias. By default, both Pyseer and our pipeline set *α* = 0.01. Although this value is close to that for ridge regression (*α* = 0), which retains all predictors in the model, it is large enough that only about 10% of 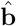 entries are non-zero.

Ten-fold (10F) and leave-one-strain-out (LOSO) [46] CV were used to assess prediction accuracy. Whereas 10F selects the training sets randomly, which can lead to instances of high similarity between test and training sequences, LOSO is a more challenging prediction task where an entire strain (= FastBAPS cluster) is predicted after training on the other strains.

### Estimation of heritability

Genetic effects at different genome sites can interact (epistasis), but we restrict attention to the narrow-sense heritability *h*^2^, with 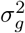 assumed to be a sum of contributions from individual sites. The LMM estimates 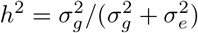 [39]. For the wg-enet heritability estimation, we used *ĥ* = *R*^2^, the proportion of phenotype variance explained by the model with *α* = 0 (ridge regression) [46].

We also estimate *h*^2^ using a modification of LDSC [47]:

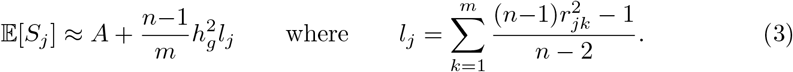

Here, *S*_*j*_ is the association test statistic at variant *j*, and *r*_*jk*_ is the sample correlation of frequency-based allele codes at variants *j* and *k* (or gene counts for the accessory genome). Following [48], prior to computing pairwise Pearson correlation coefficients we further transformed the allele codes using Gaussian quantile normalisation.

The score *l*_*j*_ involves a sum over the whole genome. In human genetics applications only a neighbourhood of *j* is included, but the presence of genome-wide LD in *S. pneumoniae* makes it difficult to define a suitable neighbourhood. The definition of *l*_*j*_ also incorporates a bias adjustment [47] that can lead to *l*_*j*_ < 0, but typically *l*_*j*_ » 1. To account for heteroskedasticity and correlations among the *S*_*j*_, the least-squares estimation of *A* and 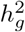 in (3) used weights 1*/*max(1, *l*_*j*_).

When choosing the testing method to generate the *S*_*j*_ for LDSC, we found that the very strong population structure effects distort the LDSC regression relationship in the absence of any adjustment, yet a fully effective adjustment for population structure was also unsatisfactory because it removed informative signal. The best compromise that we could identify between inadequate control for population structure effects and loss of association signal with effective control, was to compute the MA test statistic *S*_*j*_ in the fixed effect model (FEM):

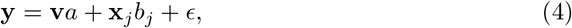

where **v** is the first principal component (PC) of the sequence distances (explaining *>* 90% of genetic variation) and *a* is the corresponding effect size. For the CD analyses, we also included the previous carriage covariate in (4). We note again that **v** does not remove all population structure effects and the *S*_*j*_ tend to be inflated, but this is not important for LDSC estimation of 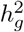 which uses the slope of the relationship of *l*_*j*_ with *S*_*j*_. Because of inadequate control of population structure using all approaches that we attempted, which included FastBAPS cluster membership indicators and additional principal components (PC), we do not report association results based on this FEM and only use the *S*_*j*_ obtained under this model within LDSC.

As well as estimating genome-wide 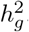, LDSC is useful for estimating the contributions to 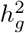 from specified genome regions. This is challenging because simply omitting variants from a heritability analysis may not exclude their effects due to strong and long-range LD. For the MIC phenotypes, we computed *ĥ* in (3) omitting effects from a known drug resistance genome region that includes the important penicillin-binding genes *pbp1a* and *pbp2x*. We first identified a set of large effect-size variants with basepair positions between 285 000 and 340 000 by clumping the frequency-coded variants using correlation threshold 0.85. These variants were used as fixed covariates when re-calculating the *S*_*j*_ for this analysis, which prevents tagging of effects from the omitted region.

### Simulation-based validation of analyses

#### Association testing

Based on the CD dataset (1 047 isolates, 134 383 variants), continuous traits were simulated under an additive model with *h*^2^ ∈ {0.1, 0.2, …, 0.5}. In each simulation, 5, 10, 15, 20, or 25 causal variants were randomly selected such that MAF *>* 0.05 and *r*^2^ < 0.2 for all pairs of causal variants. Four replicates were preformed for each of the 25 combinations of causal loci and *h*^2^, and the resulting 100 simulated datasets included a total of 1 500 causal loci (≈ 0.011% positives). Association testing was performed using the gap/SNP, major allele (MA), and treeWAS tests. For the gap/SNP test we considered both max and combi statistics.

#### Heritability estimation

We used BacGWASim [40] to simulate 1K bacterial genomes of length 250 Kb under each of two LD scenarios: lateral gene transfer rate (lgtRate) = 0.2 (Low-LD) and = 0.1 (High-LD). For each scenario and each *h*^2^ ∈ {0.1, 0.2, …, 0.9}, we simulated 100 continuous traits using 10 randomly-selected causal variants with MAF *>* 0.05 and *r*^2^ < 0.2. We then computed *ĥ*^2^ for each of the 1 800 traits using LMM, wg-enet and LDSC.

## Code and data availability

Code is available at https://github.com/Sudaraka88/bacterial-heritability and access details for the genetic data are provided in S1 File.

## Results

### Simulation analyses

For the gap/SNP test, the max statistic performed better than combi (Fig. 3). Only the max statistic is used in the real-data analyses below. The gap/SNP test is superior to both the MA and treeWAS tests (AUC for gap/SNP-max: 0.91 ± 0.01; gap/SNP-combi: 0.89 ± 0.01; MA: 0.85 ± 0.01 and treeWAS: 0.81 ± 0.02). At a Bonferroni corrected threshold of 0.05, the sensitivity and specificity were 0.433 and 0.986 for gap/SNP-max, 0.374 and 0.989 for gap/SNP-combi, 0.334 and 0.988 for MA and 0.238 and 0.996 for treeWAS.

**Fig 3.**
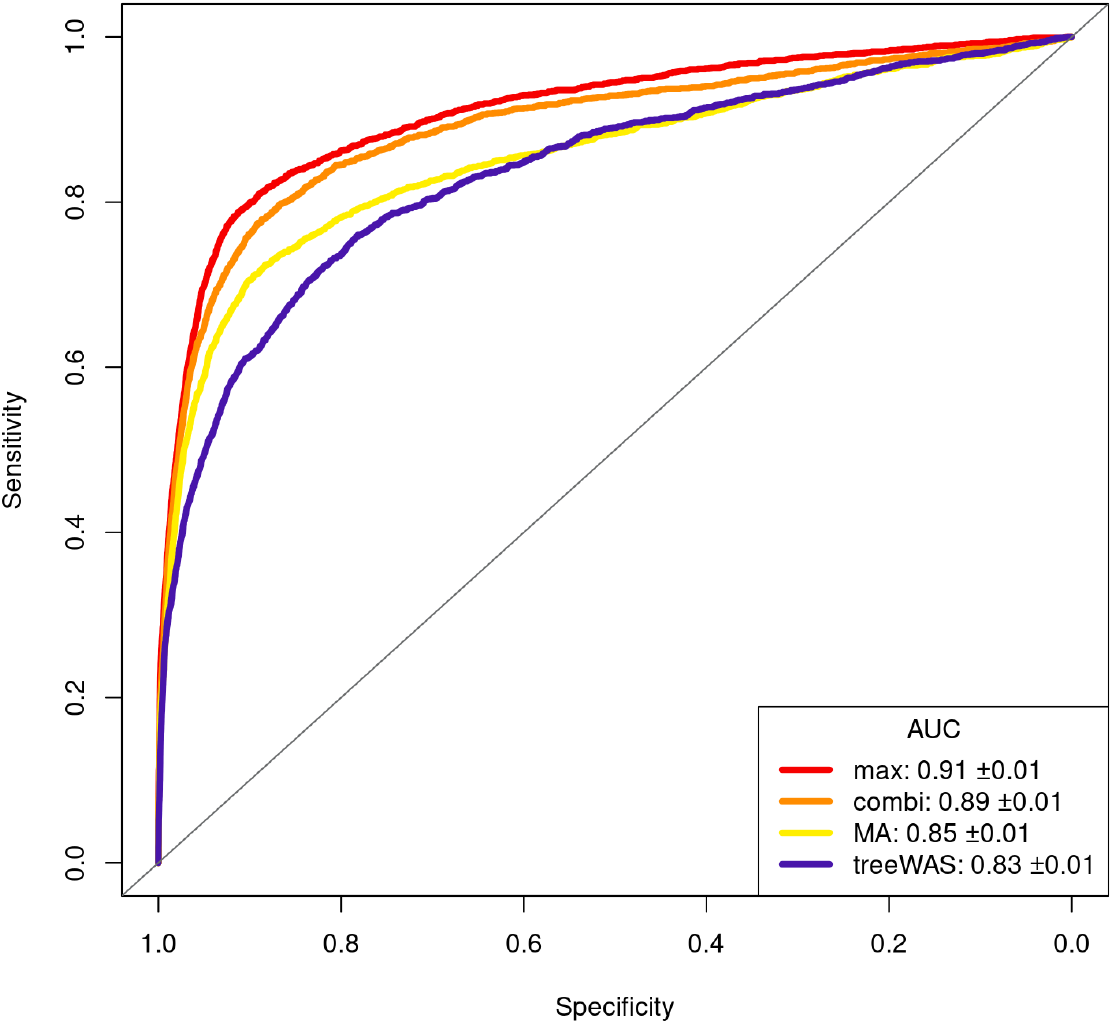
ROC curves for association tests. The gap/SNP test with max statistic outperformed the major-allele (MA) and treeWAS tests (see legend box) in our simulation study which used simulated traits and sequences from the CD dataset.

In heritability estimation, LDSC is the best-performing method, although it tends to slightly under-estimate, particularly in the high-LD scenario and for higher *h*^2^ (Fig. 4). LMM greatly over-estimates, particularly in the range 0.2 < *h*^2^ < 0.6, and wg-enet also tends to generally over-estimate. Comparatively, the bias in LDSC is quite negligible. However, wg-enet does slightly better than LDSC in the high-LD setting for *h*^2^ *>* 0.8. Overall, both LMM and wg-enet estimates are more precise than LDSC, but less accurate.

**Fig 4.**
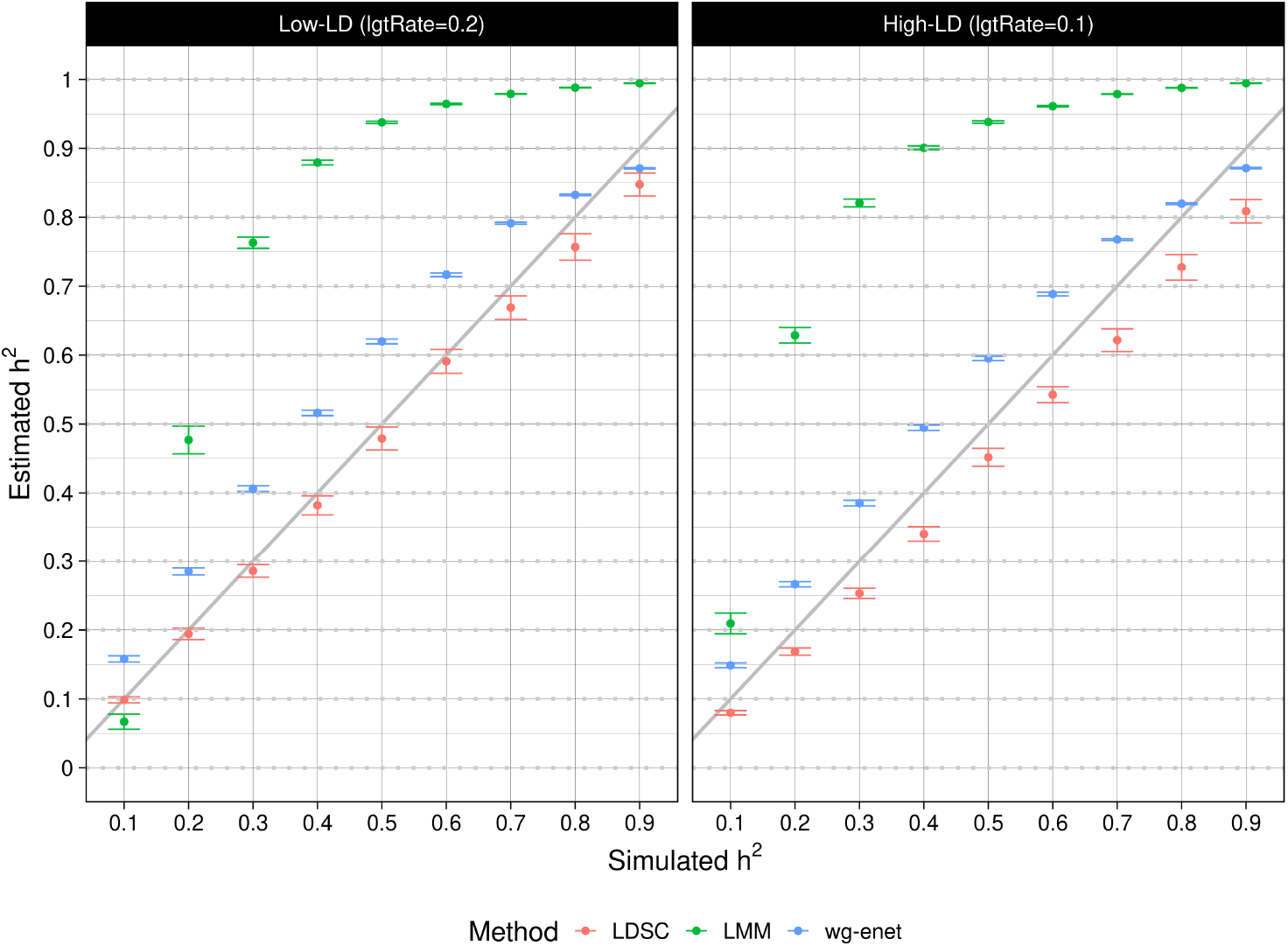
Estimating the heritability of simulated bacterial phenotypes. In the low-LD genome simulation (lateral gene transfer rate (lgtRate) = 0.2), average relative errors for LMM (green), wg-enet (blue) and LDSC (red) are 28.3±0.6%, 7.7±0.2% and −2.1±0.4%. In the high-LD genome simulation (lgtRate = 0.1), average errors for LMM (green), wg-enet (blue) and LDSC (red) 32.4±0.6%, 6.0±0.2% and −5.6±0.4%. The error bars show estimated standard error of the mean.

### Carriage duration (CD)

None of the 2 310 tested accessory genes were associated with CD. Similarly there were no genome-wide significant results among the 44 097 gap and 91 822 SNP tests at core genome variants (Fig. 5). The shared-host random effect explained 1.4% of variance for CD, and *R*^2^ = 0.0022 for the previous carriage fixed effect (*β* = −0.097, SE = 0.026).

**Fig 5.**
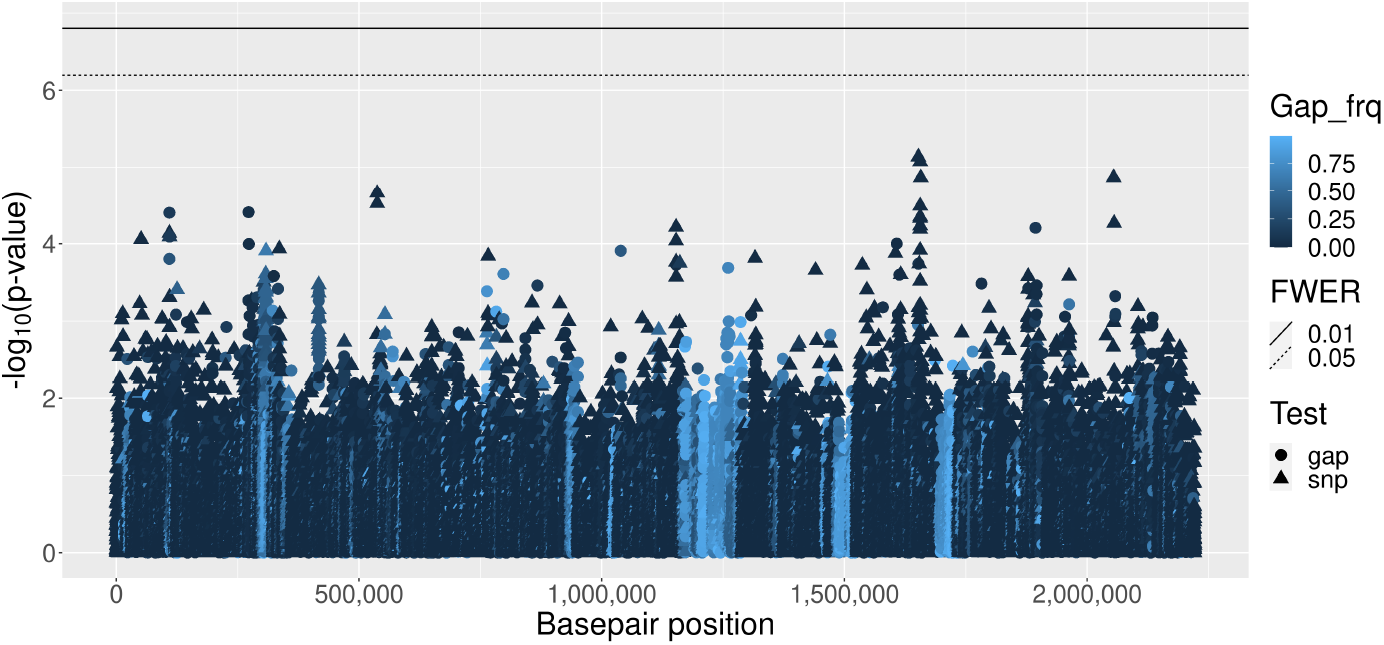
Carriage duration (CD) Manhattan plot for core genome variants. Accessory genes are not shown. See legend for shading that indicates gap frequency and symbol shape indicating gap or SNP test. Basepair positions are obtained from the ATCC700669 reference genome alignment.

The QQ-plot (S3 Fig) indicates some inflation of test statistics suggestive of population structure effects (genome inflation factor, GIF = 1.44). The MA test also identified no associations (GIF = 1.22, S4 Fig) and treeWAS identified 3 hits in 2 genes: *purF* and *polA* (S5 Fig).

Despite the lack of associations for CD, prediction accuracy (Table 1) and heritability estimates (Table 2) are significantly above zero, suggesting a polygenic trait. As expected, LOSO prediction is less accurate than 10F CV. Pangenome estimates from wg-enet, LMM and LDSC are similar (0.32 ≤ *ĥ*^2^ ≤ 0.34) with all methods also agreeing on a negligible contribution to *h*^2^ from the accessory genome. LDSC analyses also confirmed only a small contribution to *h*^2^ from the known drug-resistance region (see S6 Fig for LDSC plots). Furthermore, phenotype prediction with allele frequency-based coding of variants slightly outperformed MA coding (S2 Appendix and S7 Fig).

**Table 1.** Phenotype prediction. Mean squared error (MSE) and the correlation between observed and predicted test values using 10-fold (10F) and leave-one-strain-out (LOSO) cross validation (CV). Predictions were performed using a wg-enet model (*α* = 0.01) in glmnet, with frequency-based allele coding (all five alleles coded according to their frequency). Approximately 2% of available predictors were used for CD and 1% were used for the two MIC phenotypes. For corresponding results from MA coded variants, see S2 Appendix.

| Phenotype<br>(log scale) | 10F CV |  | LOSO CV |  |
| --- | --- | --- | --- | --- |
|  | MSE (SE) | Cor (SE) | MSE (SE) | Cor (SE) |
| CD | 0.10 (0.004) | 0.55 (0.022) | 0.12 (0.005) | 0.44 (0.025) |
| Ceftriaxone MIC | 0.03 (0.002) | 0.91 (0.005) | 0.08 (0.003) | 0.77 (0.005) |
| Penicillin MIC | 0.04 (0.003) | 0.91 (0.005) | 0.13 (0.051) | 0.69 (0.014) |

**Table 2.** Heritability estimates (*ĥ*^2^). The upper and lower values in each cell are for core genome and pangenome (= core genome plus accessory genes). Under “w/o DR” are results from analyses that omit effects from a genome region that is known to be associated with drug resistance.

| Phenotype | LDSC |  |  |  |
| --- | --- | --- | --- | --- |
|  | wg | w/o DR | enet | LMM |
| CD | 0.34 | 0.30 | 0.34 | 0.32 |
| *with* accessory genes | 0.34 | 0.31 | 0.34 | 0.32 |
| Ceftriaxone MIC | 0.86 | 0.22 | 0.92 | 0.98 |
| *with* accessory genes | 0.87 | 0.22 | 0.93 | 0.98 |
| Penicillin MIC | 0.72 | 0.40 | 0.94 | 0.98 |
| *with* accessory genes | 0.72 | 0.41 | 0.94 | 0.98 |

We also performed association testing on all 1 612 isolates linked to a carriage episode. This analysis identified four sites at basepair positions 1 522 542–1 522 896, near the previously-reported phage hit based on *k*-mer analysis [46]. However, our 4 hits are due to the same 15 isolates, of which 6 are from the same long (517 day) episode (see detailed results in S1 Appendix). Furthermore, when the all-isolates dataset was analysed using treeWAS, 9 associations were identified (see S3 Appendix), but these did not include *purF* and *polA* (reported above) nor the region identified in our LMM analyses. We conclude that we are unable to reliably identify individual associations for CD, but there is good evidence for it being a moderately-heritable polygenic trait.

### Minimum inhibitory concentration (MIC) phenotypes

For both MIC phenotypes, from the 2 242 accessory genes tested, one (with Panaroo label group 102) showed genome-wide significant association. Gap and SNP tests were performed at 36 020 and 97 224 core genome sites, respectively. For ceftriaxone MIC and penicillin MIC, respectively, 998 and 833 variants showed genome-wide significance (Figs 6, 7), and 688 and 504 of these were within annotated gene regions of the ATCC700669 reference genome [49] (Table 3). Approximately 35% of hits were from the gap test, associations that have largely been ignored in previous analyses. For ceftriaxone MIC and penicillin MIC, GIF = 1.14 and 1.28 respectively, but the QQ plots (S9 Fig) suggest that, rather than genome-wide inflation caused by population structure, GIF *>* 1 is due to a large fraction of the genome showing causal association with these highly-heritable, polygenic traits.

**Fig 6.**
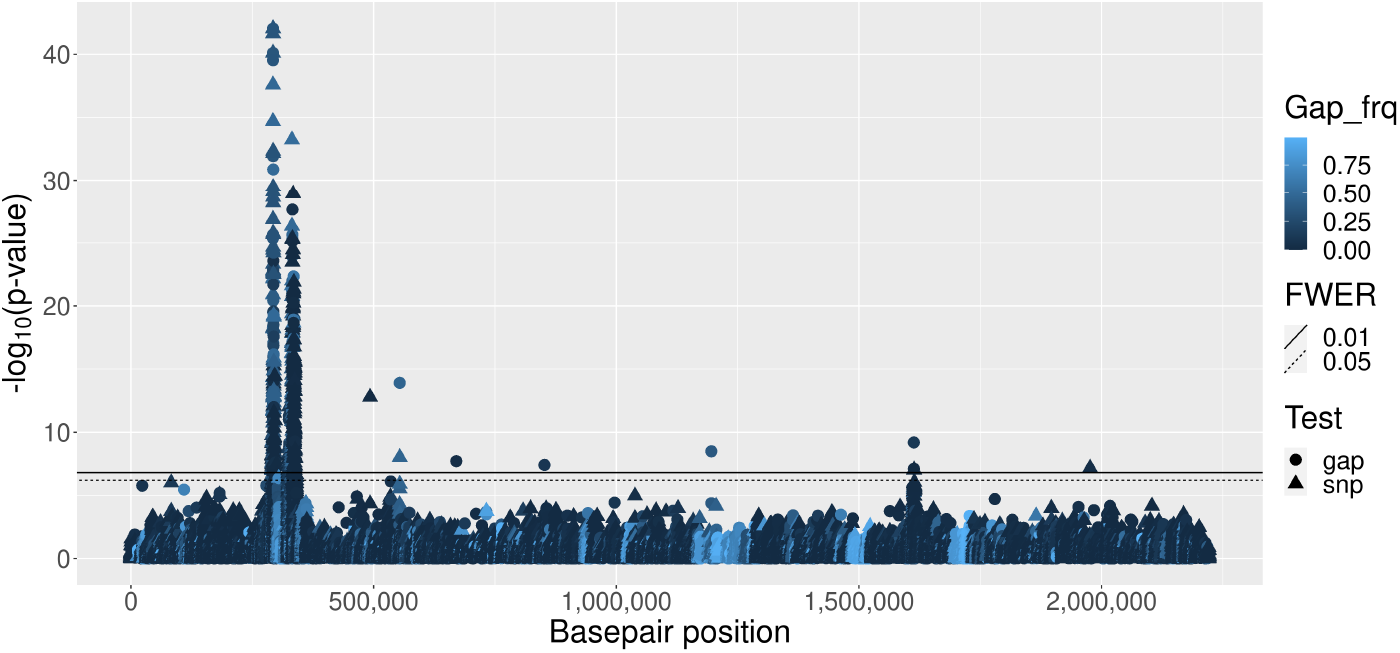
Ceftriaxone MIC Manhattan plot. The shading and symbol shapes (see legend) are the same as for Fig. 5

**Fig 7.**
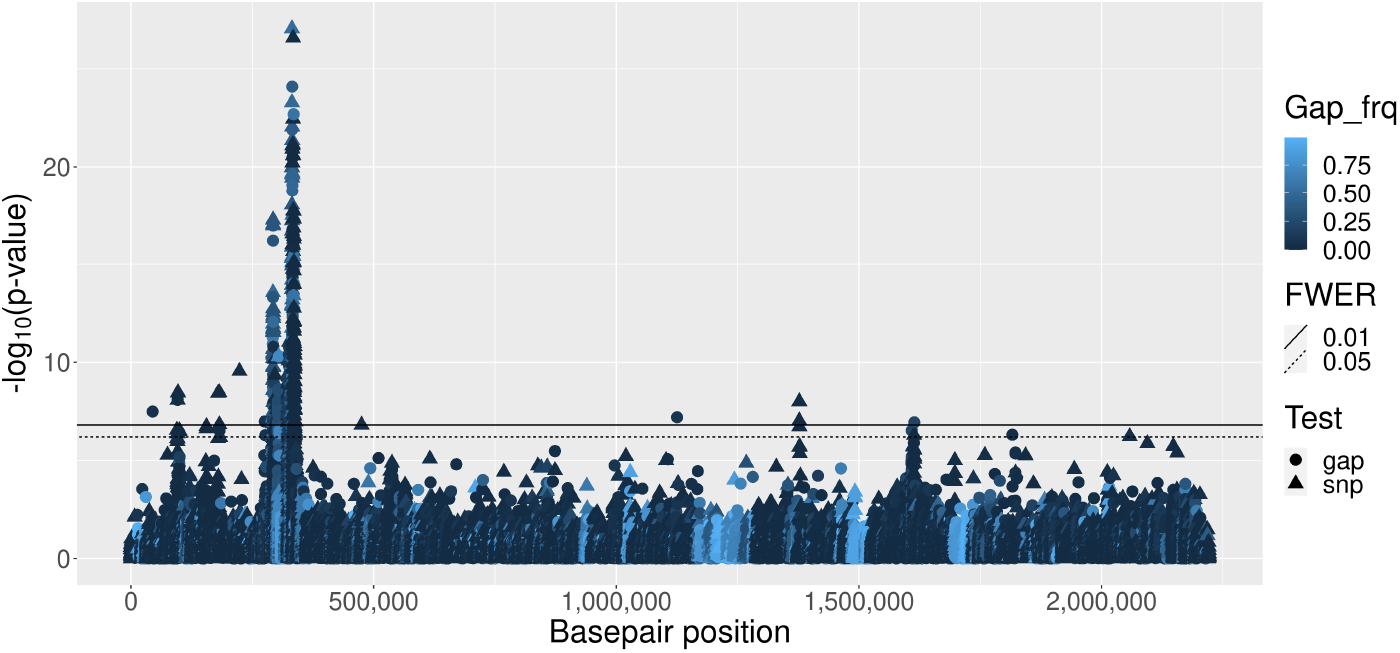
Penicillin MIC Manhattan plot. See Fig. 6 caption for details.

**Table 3.** Genes showing significant association with MIC phenotypes.

| Phenotype (log) | Core genes | Acc. gene |
| --- | --- | --- |
| Ceftriaxone only | mraW, clpL, csrR, rplK, aliB, plr, valS |  |
| Both MIC phenotypes | pbp1a, aliA, pbp2x, mraY, recU, gnd, dexB, luxS, wzg, pbp2b | group_102 |
| Penicillin only | aliB, clpL, wzd, wzh, blpY, galK, hasC, leuB, leuS, murF, recO |  |

For ceftriaxone MIC, the largest statistics are of similar magnitude for gap and SNP tests (Fig. 8), but for low allele frequencies there are few large gap statistics and many large SNP statistics, suggesting that there are few rare deletions, but many rare nucleotides of large effect. There are also few large gap statistics with frequency *>* 0.6, suggesting few sequence insertions of large effect. Many large SNP statistics with frequency above 0.4 were not recorded as significant under the MA test, which may reflect a benefit of frequency-based allele coding.

**Fig 8.**
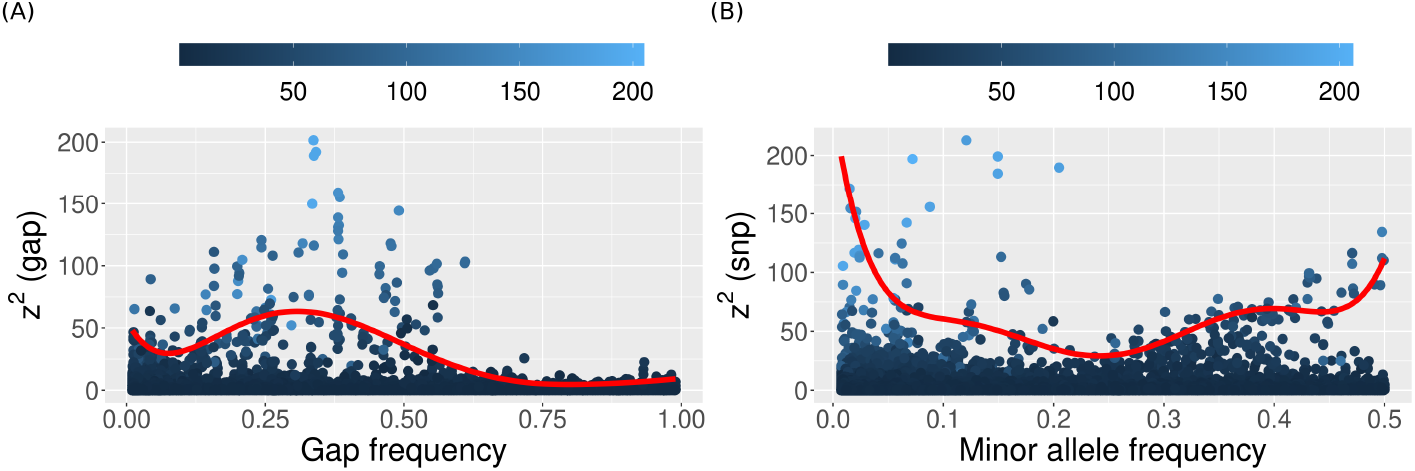
Association test statistics against variant frequency for ceftriaxone MIC. Each point shows the *z*^2^ statistic from (A) gap and (B) SNP test at a core genome variant. The *x*-axis shows frequencies of (A) gap and (B) minor nucleotide as a fraction of all nucleotides. Points are shaded according to the major-allele (MA) test statistic and the red curve shows 7^th^ order regression fit for the 90^th^ percentile [50]. See S8 Fig for this analysis on the other two phenotypes.

Consistent with the simulation results, the gap/SNP test performed better than the MA and treeWAS tests, identifying more associations (1 831 vs 1 419 vs 206) with lower GIF (1.14 vs 1.20 and 1.28 vs 1.56). For further details from (1) the major allele test, see S10 Fig and S11 Fig and (2) treeWAS analysis, see S12 Fig, S13 Fig. The list of genes identified from both analyses are in S3 Appendix.

As expected from the large number of associations, prediction accuracy for both MIC phenotypes is very high under 10F CV (Table 1), but less so for LOSO CV, with high SE for penicillin MIC indicating hard-to-predict clusters (S14 Fig).

The values of *ĥ* also reflect the simulation studies, with LMM *>* wg-enet *>* LDSC for both MIC phenotypes (Table 2). Whereas LMM and wg-enet agreed closely across the two MIC phenotypes, the LDSC *ĥ* differ substantially, consistent with the higher LOSO prediction accuracy and higher numbers and significance of associations for ceftriaxone MIC compared with penicillin MIC (see S6 Fig for LDSC plots). Using LDSC we also estimate that around a quarter of *h*^2^ for ceftriaxone MIC can be attributed to the known drug resistance region, which represents only 2.5% of the core genome. This fraction rises to just over half of *h*^2^ for penicillin MIC.

## Discussion

We have identified several improvements over previous approaches for association, prediction and heritability analyses for quantitative bacterial traits, verified through simulation studies and real-data analyses. We present the first genomic analyses of *S. pneumoniae* minimum inhibitory concentration (MIC) for the beta-lactam antibiotics ceftriaxone and penicillin, finding many novel associations and high heritability. Prediction of MIC traits was correspondingly accurate under 10F CV.

The genome regions identified as associated with the MIC phenotypes overlap those previousy reported for the binary AMR phenotypes, even in the case of ceftriaxone for which none of the tested isolates was resistant. Many of the associated genes are in the peptidoglycan biosysthesis pathway, including penicillin binding proteins (PBPs: *pbp1a, pbp2b, pbp2x*) and transferases required for cell wall biogenesis (*mraY* and *mraW* for ceftriaxone MIC). A single heat shock protein (*clpL*) and a gene from the recombination pathway (*recU*) were also identified as associated. When present, the group 102 accessory gene is located adjacent to *pbp1a*, which generates an enzyme involved in cell wall remodelling, which may contribute to the association signal for the MIC phenotypes. However, most of the genes identified for the MIC phenotypes are in tight linkage with the three PBPs and may not represent independent effects.

We found no reliable associations for *S. pneumoniae* carriage duration (CD), but strong evidence that it is a polygenic trait of moderate heritability (*ĥ* ≈ 0.33) that is predictable from the genome sequence (0.55 and 0.44 correlation between predicted and true phenotype under 10F and LOSO CV, respectively).

The innovations in our association analysis pipeline include separate testing of gap and SNP effects, with a permutation approach to control FWER and frequency-based allele coding. This approach performed better than the alternatives of MA and treeWAS tests, detecting more associations under good control of population structure effects.

Our phenotype prediction analysis used frequency-coded variants within a glmnet-based whole genome elastic net model.

A previous analysis of CD using data from the same study [3], provided a lower-bound *h*^2^ estimate of 0.45 using warped-lmm [51], concluding that CD is a highly heritable trait. Our estimates are lower, which may be due to our decision to use only one isolate per CD episode (S1 Appendix).

Penicillin AMR *ĥ* ^2^ in the Maela data set was recently reported in the range 0.67–0.83 [4]. Our most reliable (LDSC) estimate for the quantitative penicillin MIC phenotype is within this range (Table 2). For ceftriaxone MIC, *ĥ* is even higher, consistent with the good prediction performance.

The reduction in *h*^2^ for penicillin MIC by more than half on removing known drug resistance genome regions in *S. pneumoniae* contrasts with results from *M. tuberculosis*, where the largest reduction in *h*^2^ (measured using GEMMA [52]) was only 27% [9], which is close to our result for ceftriaxone MIC.

Our results support the use of linear mixed models for association analysis, with separate testing of gap and SNP effects, the latter using frequency-based allele coding. We also support the use of wg-enet for prediction of quantitative traits and we find that LDSC performs best for heritability analyses. Further work is required to assess optimal strategies in a wider range of settings for population structure in bacterial genomes.

## Supporting information

**S1 Appendix. Results from the carriage duration analysis using the dataset comprising all** 1 612 **isolates sampled during a positive episode**.

(PDF)

**S2 Appendix. Phenotype prediction using MA frequency coded variants**.

(PDF)

**S3 Appendix. Genes identified with MA tests (LMM and treeWAS)**.

(PDF)

**S1 Fig. Phenotype distribution**. Top and bottom rows show the distribution of the three phenotypes before and after log_10_ transformation.

**S2 Fig. Phylogenetic trees from the ClonalFrameML analysis**. Mid-point rooted, ‘recombination-aware’ tree structure for (A) 1 047 isolates with carriage duration phenotype (measured in days and indicated by tip colour) and (B) 1 332 isolates with MIC phenotype (measured in μg ml^*−*1^ and tip colour indicates the distribution of penicillin MIC).

(PDF)

**S3 Fig. QQ plot for carriage duration from the GAP/SNP analysis**.

(PDF)

**S4 Fig. Manhattan plot from MA tests of association with CD**. Testing was performed with (A) LMM and (B) FEM models. LMM did not identify any significant associations, whereas FEM identified 92 associations with GIF = 2.53, indicating genome-wide inflation due to unsatisfactory control of population structure. In FEM, population structure correction was performed using FastBAPS cluster indicator covariates. Point colour indicates the gap frequency at each site and the horizontal lines indicate Bonferroni corrected significance thresholds.

(PDF)

**S5 Fig. treeWAS analysis for CD**. Manhattan plots for (top) Terminal (middle) Simultaneous and (bottom) Subsequent scores are shown, where three hits are identified from the simultaneous test.

(PDF)

**S6 Fig. LDSC analyses for all phenotypes**. LDSC plots for (A) CD, (B) ceftriaxone MIC and (C) penicillin MIC. In each figure, subplots correspond to the **a**. core genome **b**. pangenome **c**. core genome w/o DR and **d**. pangenome w/o DR analyses and the *ĥ* are reported in Table 2.

(PDF)

**S7 Fig. Prediction accuracy with MA and frequency coding**. Allele frequency coding generally increases the correlation and reduces the mean squared error of prediction for all three phenotypes across folds and clusters. Note that the Mean squared error and correlation values here are averaged across folds and clusters, and are different from the overall accuracy results in Table 1 and S2 Appendix

(PDF)

**S8 Fig. Variation in** *z*^2^ **statistics with variant frequency** Each point shows the *z*^2^ statistic of a (A, C) gap and (B, D) SNP tested core genome variants for (A, B) carriage duration and (C, D) penicillin MIC. The *x*-axis respectively shows the gap and minor allele frequency for gap and SNP tested variants. Points are shaded according to the *z*^2^ statistic from the MA test and the red curve shows the 4^th^ order regression fit for the 90^th^ percentile of data.

(PDF)

**S9 Fig. QQ plots for MIC phenotypes from the GAP/SNP analysis**. (A) ceftriaxone MIC (B) penicillin MIC.

(PDF)

**S10 Fig. Major-allele test for ceftriaxone MIC**. Testing was performed using

(A) LMM and (B) FEM models. FEM analysis identified 13 212 hits with GIF = 16.4. See caption in S4 Fig for additional analysis and figure legend details.

(PDF)

**S11 Fig. Major-allele test for penicillin MIC**. Testing was performed using (A) LMM and (B) FEM models. FEM analysis identified 23 636 hits with GIF = 17.0. See caption in S4 Fig for additional analysis and figure legend details.

(PDF)

**S12 Fig. treeWAS analysis for ceftriaxone MIC**. Manhattan plots for (top) Terminal (middle) Simultaneous and (bottom) Subsequent scores.

(PDF)

**S13 Fig. treeWAS analysis for penicillin MIC**. Manhattan plots for (top) Terminal (middle) Simultaneous and (bottom) Subsequent scores.

(PDF)

**S14 Fig. Prediction performance**. Prediction performance of (A,B) carriage duration, (C,D) ceftriaxone MIC and (E,F) penicillin MIC phenotypes, assessed using (A,C,E) 10F and (B,D,F) LOSO CV. The *x* and *y* axes denote the true and predicted values, respectively and point colour represents the fold or FastBAPS cluster. Mean squared error and correlation values in Table 1 and S2 Appendix are computed using all values shown here.

(PDF)

**S1 File. Metadata for *S. penumoniae* isolate reads used in this study**. (CSV)

## Acknowledgments

SM and DB were supported by grant DP190103188 from the Australian Research Council. JC was funded by the ERC grant no. 742158.

