## Supplementary material for "Genome-wide association, prediction and heritability in bacteria": S1_Appendix

### S1 Appendix. Results from the carriage duration analysis using the dataset comprising all 1 612 isolates sampled during a positive episode

We performed association testing on the dataset comprising all isolates (1 612) that were linked to a carriage episode. An additional 733 accessory genes were tested here, none of which showed association, and Gap and SNP tests were performed on 77 614 and 115 745 core genome variants in total, respectively, with 44 119 subject to both tests. Four sites (gap tested only) were significant ($p = 4 \times 10^{-9}$) at a Bonferroni corrected threshold of 0.01. These all correspond to an insertion represented in 15 sequences, at basepair positions 1 522 542, 1 522 896, 1 522 934 and 1 522 964. This region is within the MM1-type prophage of the ATCC 700669 genome. An association with CD was previously reported at a nearby locus using k-mer analysis [1]. The 15 insertion sequences have a mean CD of 308 days (SD 183), compared with 145 (SD 125) for the 1 597 gap sequences. However, the association signal largely comes from 6 isolates from the same 517-day CD episode and requires further investigation to confirm its significance. Estimates of $h^2$ (Table 1) were naturally higher for this larger dataset where 337 episodes are over-represented by isolates.

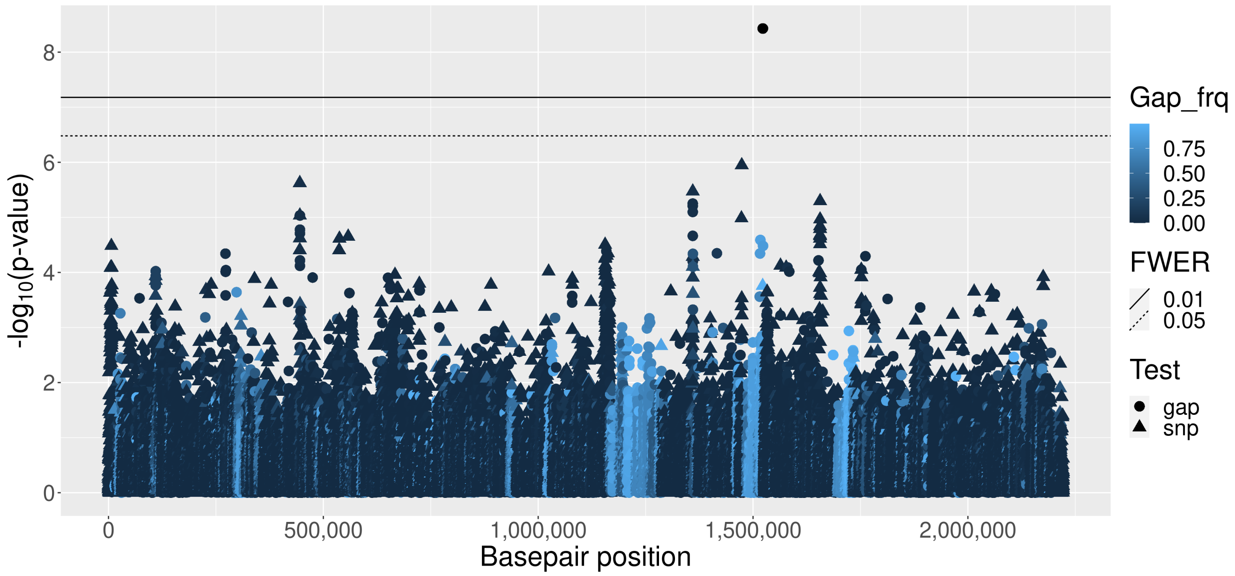


Figure 1: **Carriage duration (CD) Manhattan plot for core genome variants.** Accessory genes are not shown. See legend for shading that indicates gap frequency and symbol shape indicating whether the test of association was based on gaps or SNP variation. Basepair positions are obtained from the ATCC700669 reference genome alignment.

Table 1: **Heritability estimates ($\hat{h}^2$).** Top and bottoms rows show $\hat{h}^2$ for the core and pan genomes, respectively.

| Phenotype (log) | Pyseer | | LDSC |
|---|---|---|---|
| | LMM | wg-enet | |
| CD - all isolates | 0.61 | 0.52 | 0.52 |
| *with* Accessory genes | 0.61 | 0.58 | 0.54 |
