## Supplementary material for "Genome-wide association, prediction and heritability in bacteria": S2_Appendix

### S2 Appendix. Phenotype prediction using major allele frequency coded variants

Table 1: **Phenotype prediction with major allele coded variants.** Mean squared error (MSE) and the correlation between observed and predicted test values using 10-fold (10F) and leave-one-strain-out (LOSO) cross validation (CV). Predictions were performed using a wg-enet model in glmnet. Approximately 1.5% of available predictors were used for CD and 1% were used for the two MIC phenotypes.

| Phenotype | 10F CV | | LOSO CV | |
|---|---|---|---|---|
| (log scale) | MSE (SE) | Cor (SE) | MSE (SE) | Cor (SE) |
| CD | 0.100 (0.005) | 0.540 (0.022) | 0.113 (0.004) | 0.457 (0.024) |
| Ceftriaxone MIC | 0.031 (0.002) | 0.910 (0.005) | 0.086 (0.004) | 0.744 (0.012) |
| Penicillin MIC | 0.042 (0.003) | 0.909 (04005) | 0.120 (0.005) | 0.731 (0.013) |
