## Supplementary material for "Genome-wide association, prediction and heritability in bacteria": S3_Appendix

### S3 Appendix. Genes identified by pyseer-LMM and treeWAS analyses

The LMM MA test identified 817 core-variant associations for ceftriaxone MIC, 524 of which were in 22 genes, 13 of them also identified by the gap/SNP test. For penicillin MIC, 602 associations were identified, 444 of which were mapped to 16 genes, also 13 in common with the gap/SNP test. No accessory gene associations were identified for either MIC phenotype.

Table 1: **Associated genes identified from pyseer-LMM.** All variants above the significance threshold in any test is checked for annotations in the ATCC700669 reference genome for associated genes.

| Phenotype (log) | Core genes | Accessory genes |
|---|---|---|
| CD - 1-isolate/episode | - | - |
| CD - all isolates | - | - |
| Ceftriaxone MIC | pbpX, pbp1A, aliA, mraY, recU, mraW, gnd, clpL, penA, csrR, dexB, luxS, accD, aldB, comFC, folC, lepA, mtsC, pabB, rplK, valS, wzh | - |
| Penicillin MIC | pbp1A, aliA, pbpX, recU, mraY, wzg, dexB, gnd, luxS, clpL, csrR, leuB, leuS, potD, recO, smc | - |

TreeWAS only identified 140 and 66 core-genome associations for ceftriaxone and penicillin MIC, respectively.

Table 2: **Associated genes identified from treeWAS.** All variants above the significance threshold in any test is checked for annotations in the ATCC700669 reference genome for associated genes.

| Phenotype (log) | Core genes | Accessory genes |
|---|---|---|
| CD - 1-isolate/episode | purF, polA | - |
| CD - all isolates | dgk, lacF, msrAB, murM, lacG2, alsS, polC, hrcA, dpr | - |
| Ceftriaxone MIC | pbp1a, pbp2b, pbp2x, mraY, clpL, recU, aliA, dexB, gnd | - |
| Penicillin MIC | pbp1a, pbp2b, pbp2x, recU, mraY, clpL, luxS, gnd | - |
