## Supplementary figures and images for "Genome-wide association, prediction and heritability in bacteria"

### S1_Fig

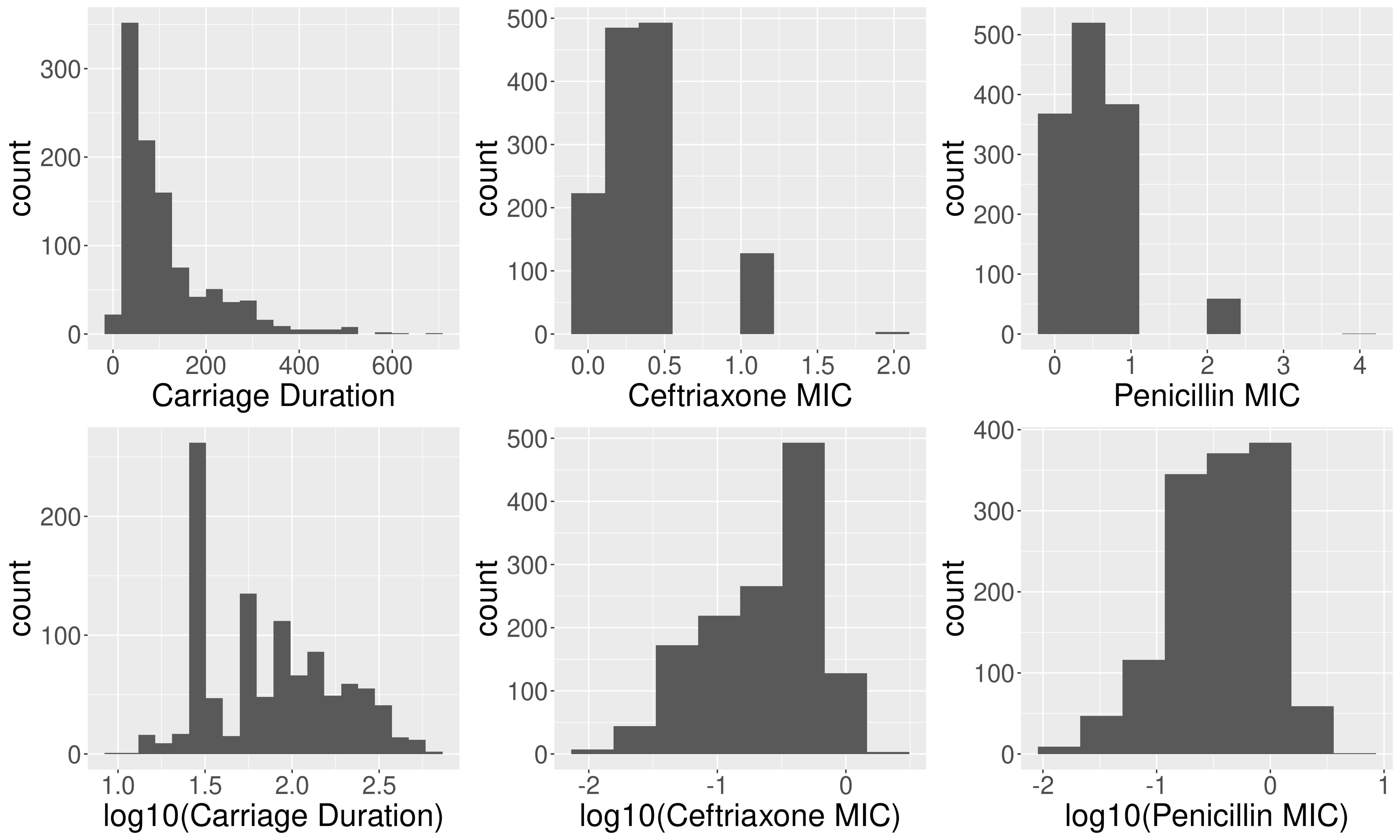

### S2_Fig

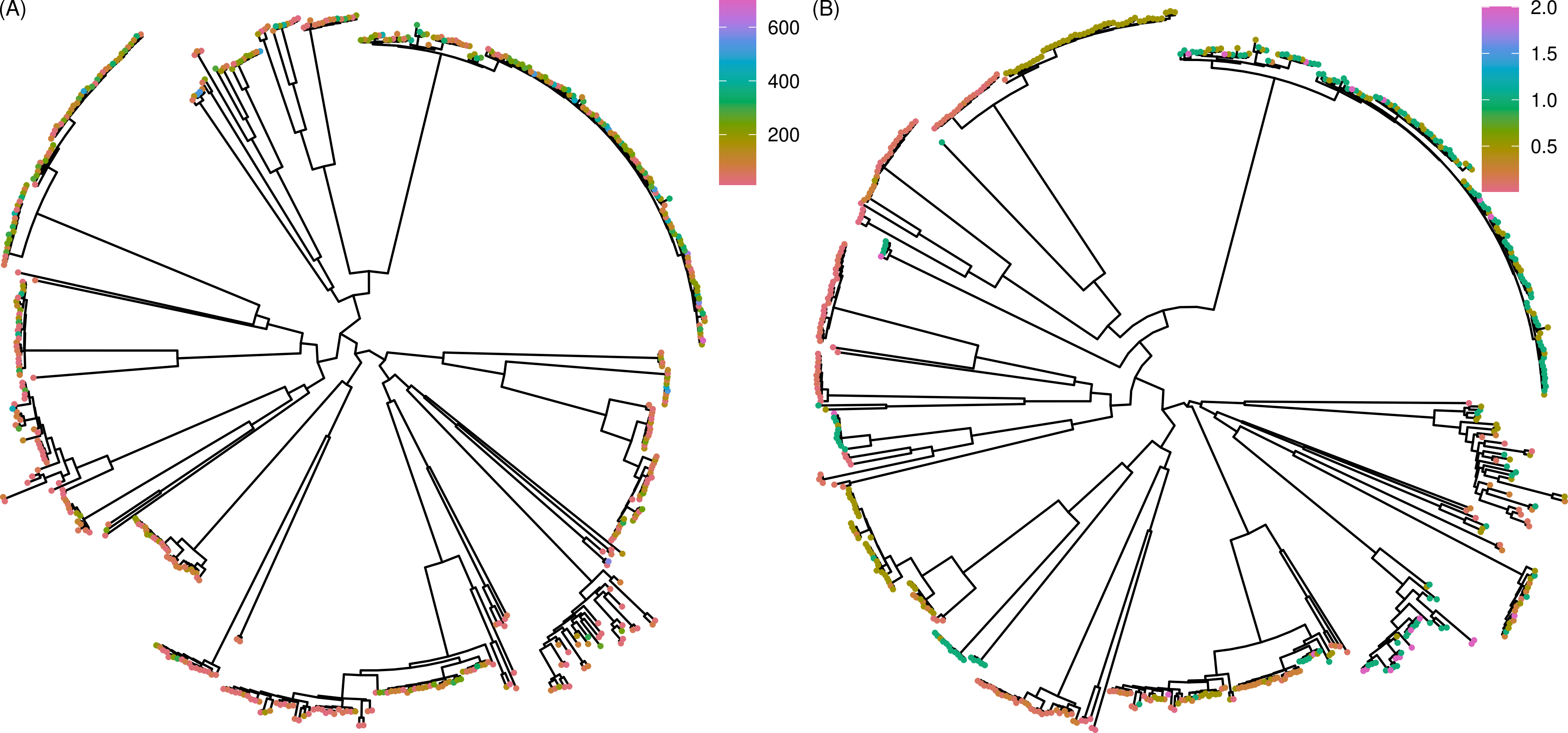

### S3_Fig

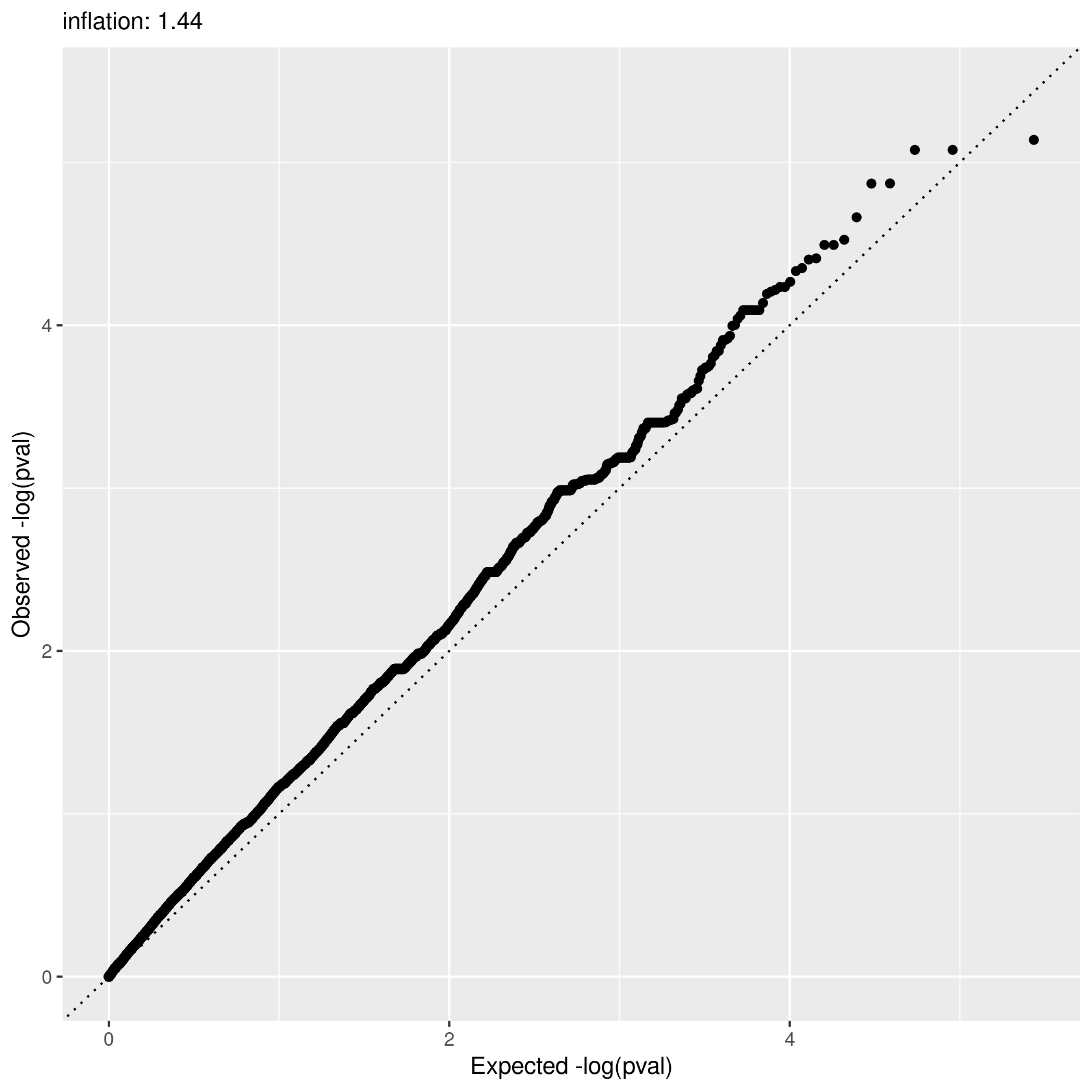

### S4_Fig

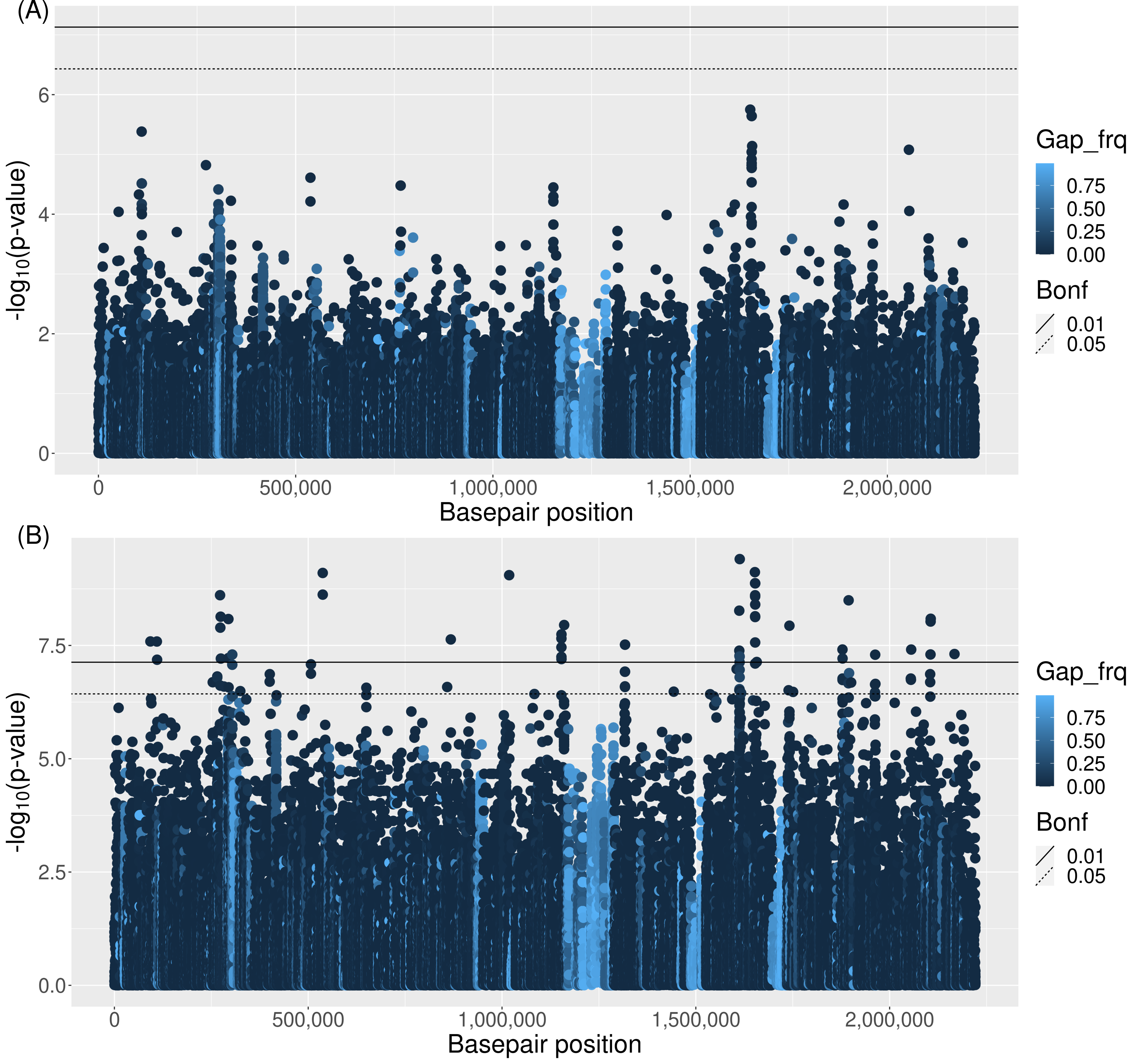

### S5_Fig

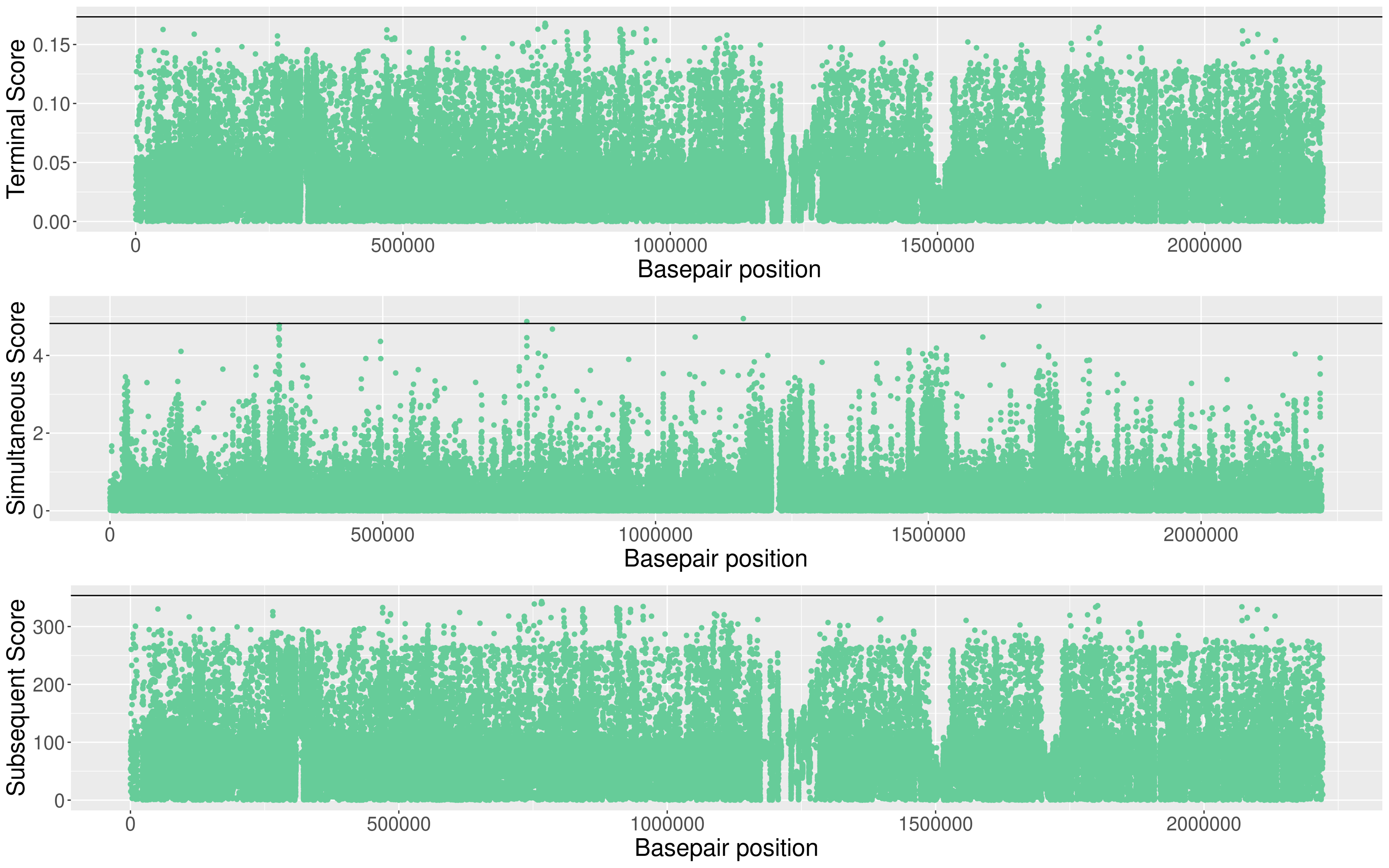

### S6_Fig

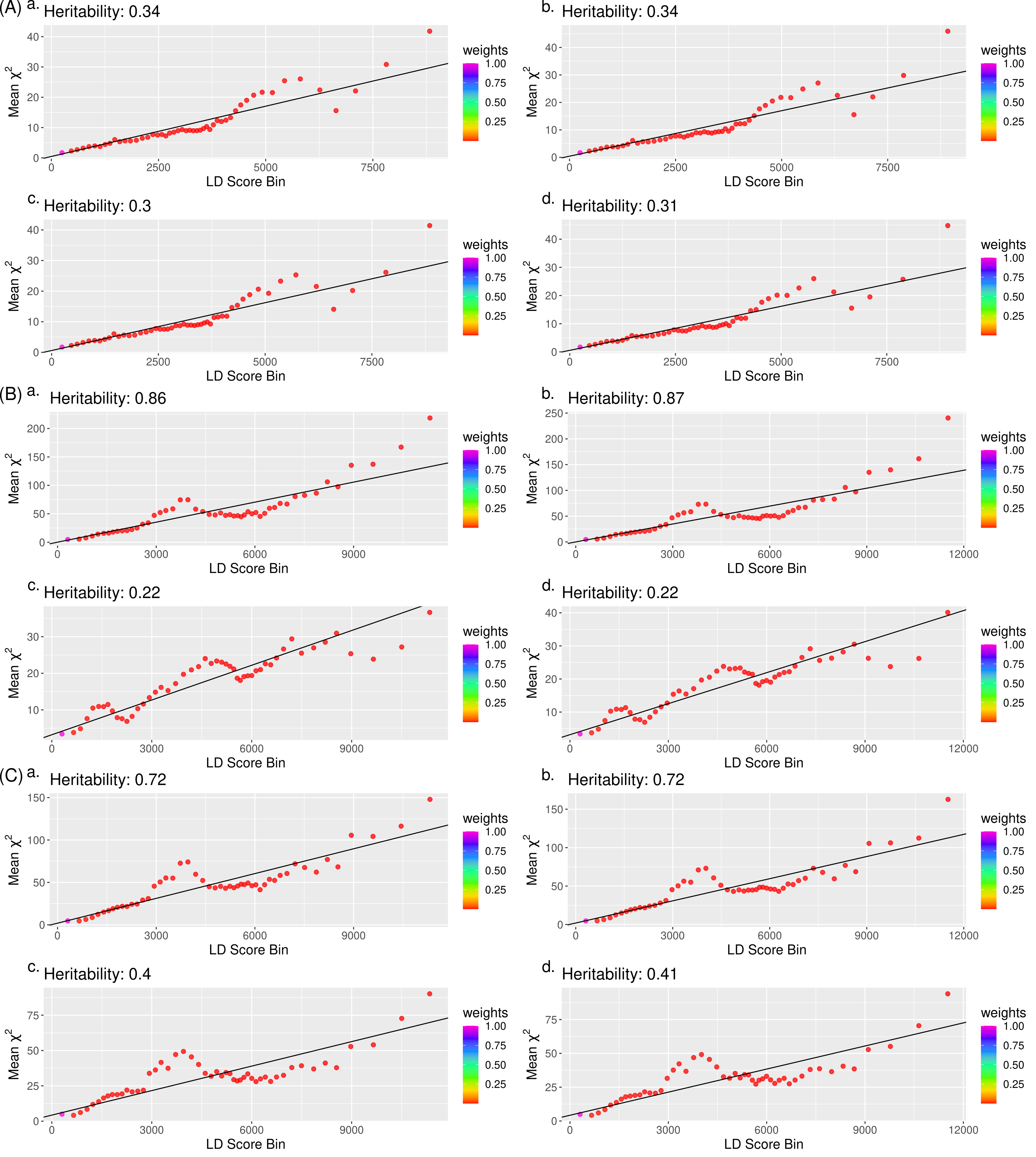

### S7_Fig

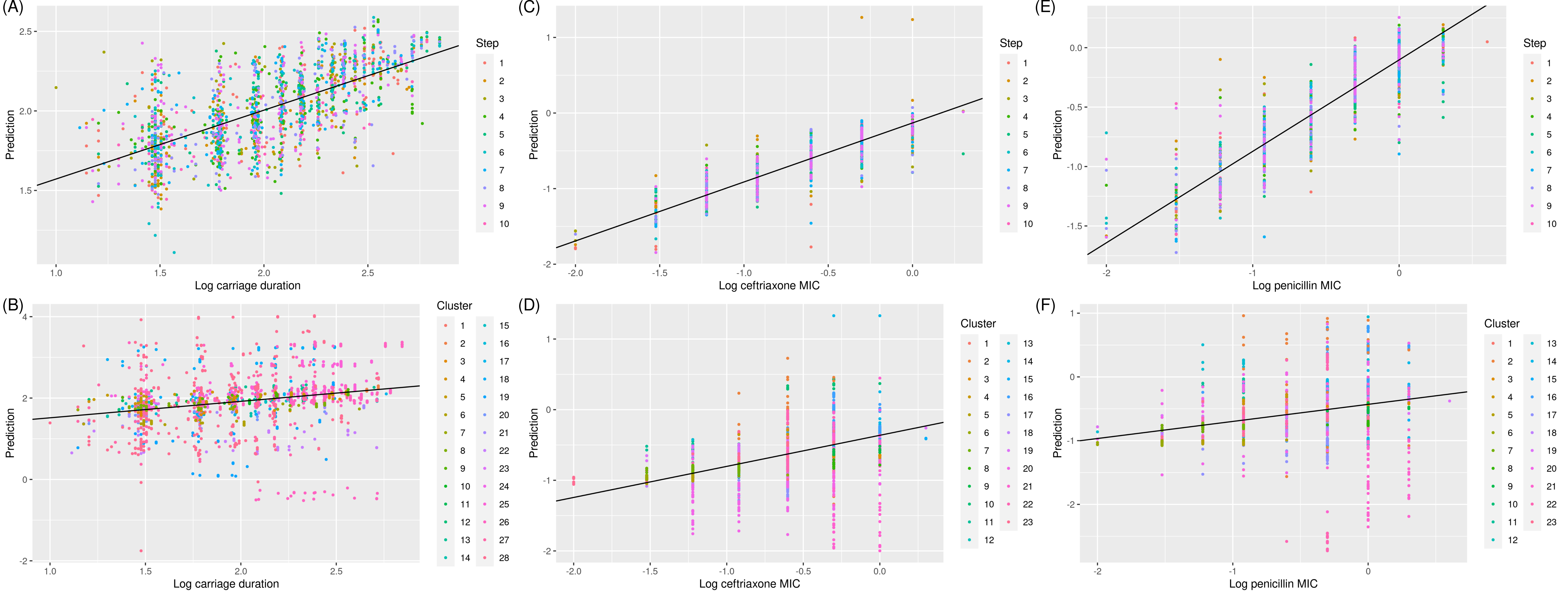

### S8_Fig

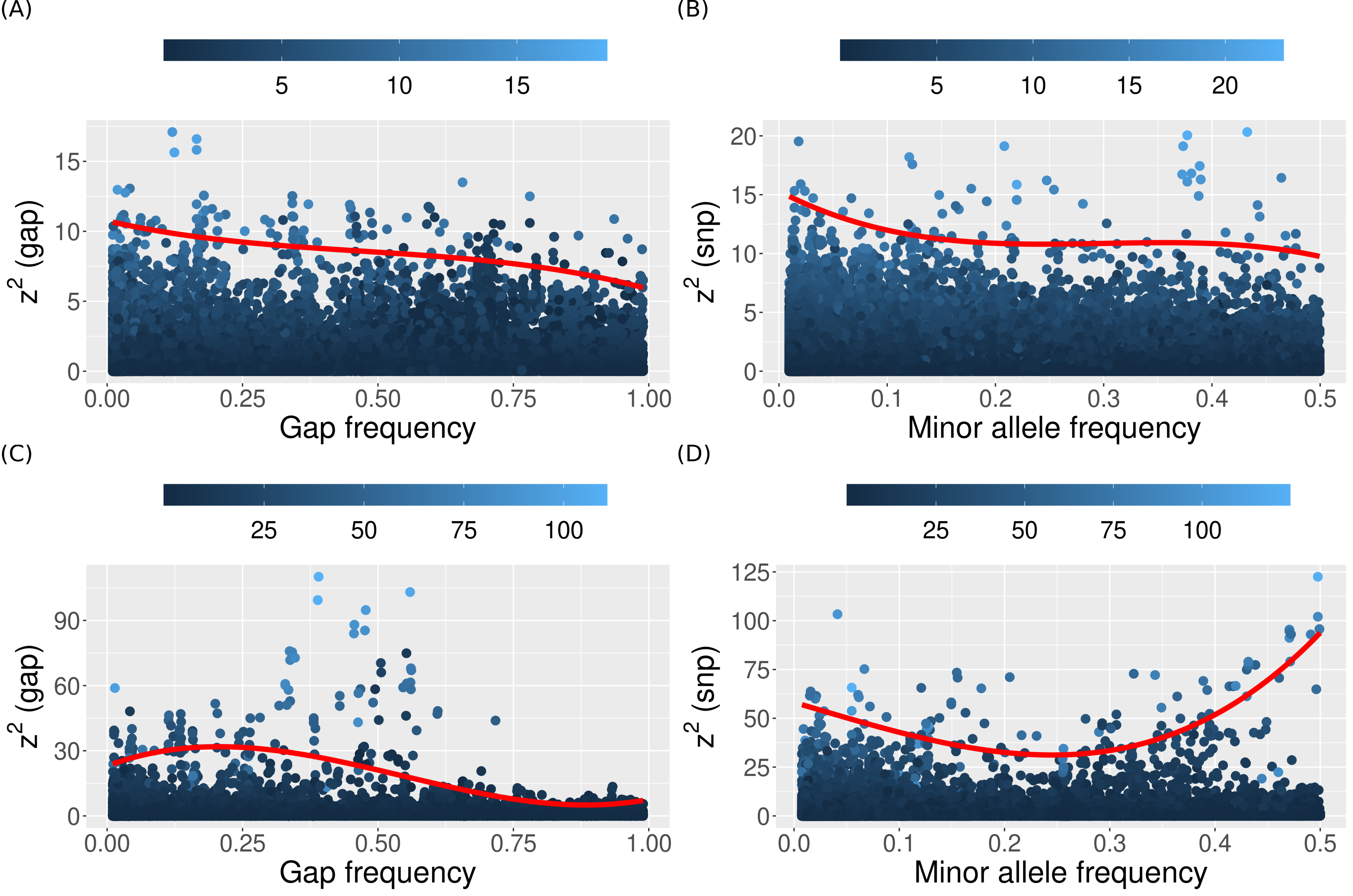

### S9_Fig

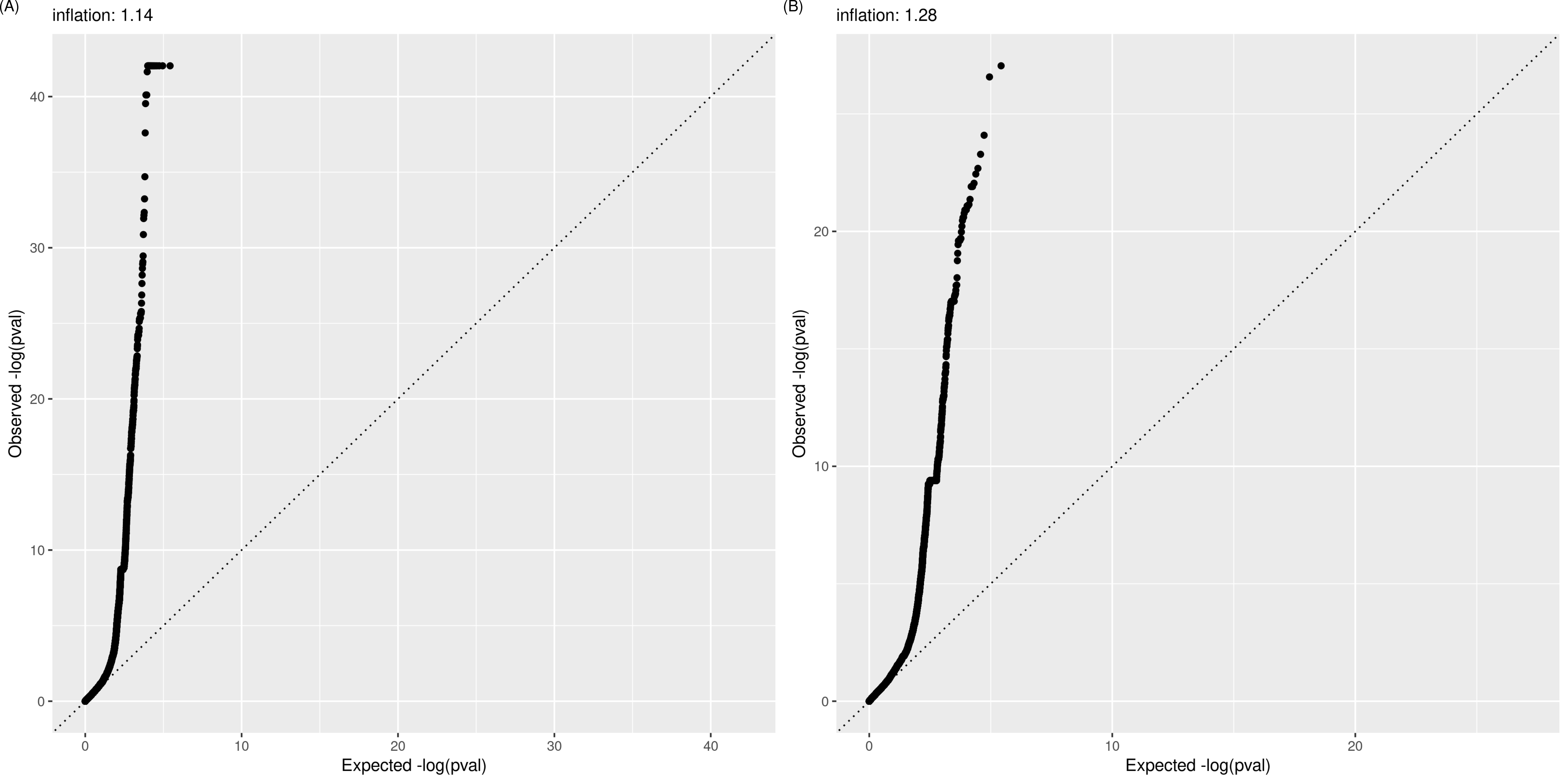

### S10_Fig

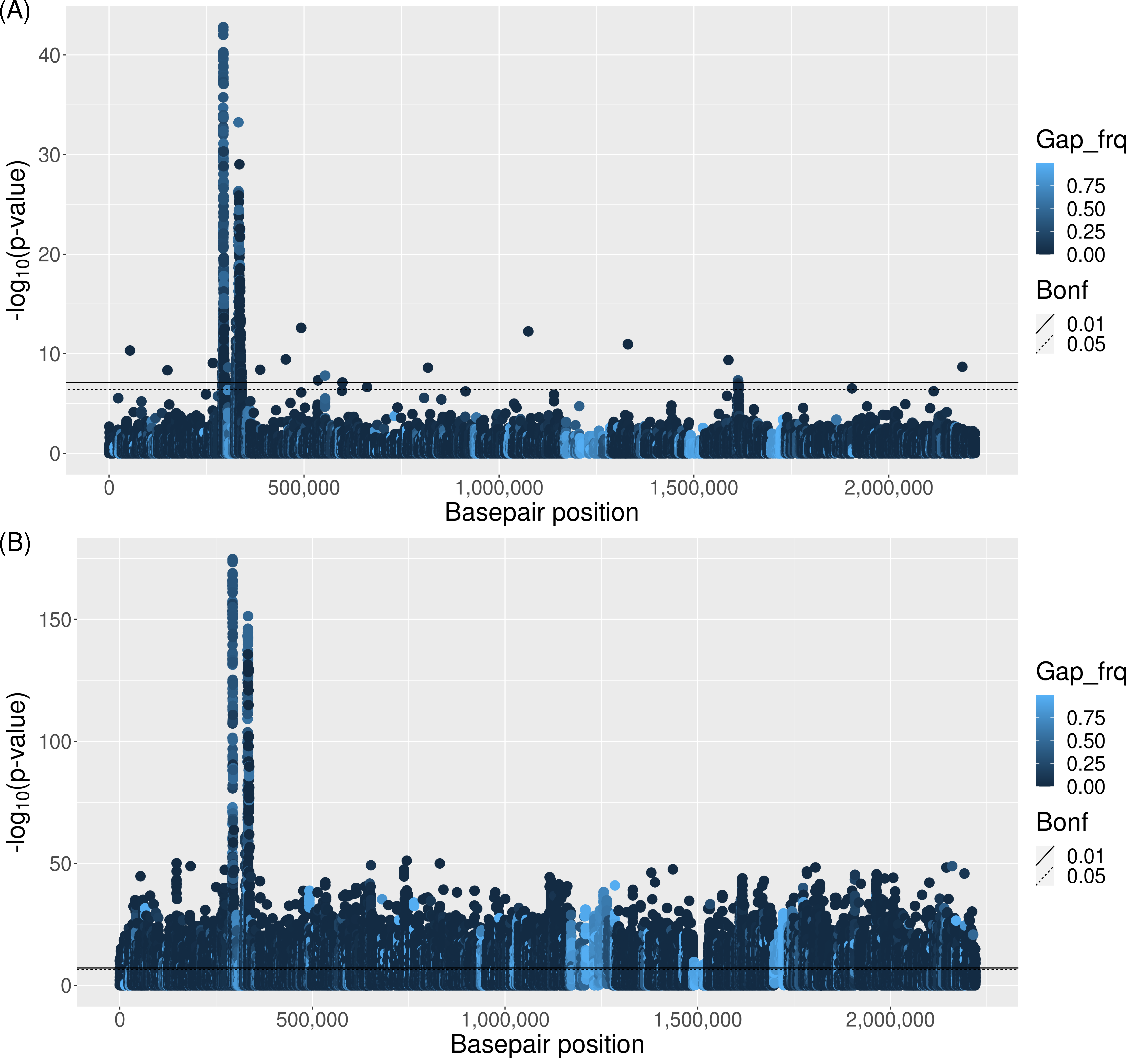

### S11_Fig

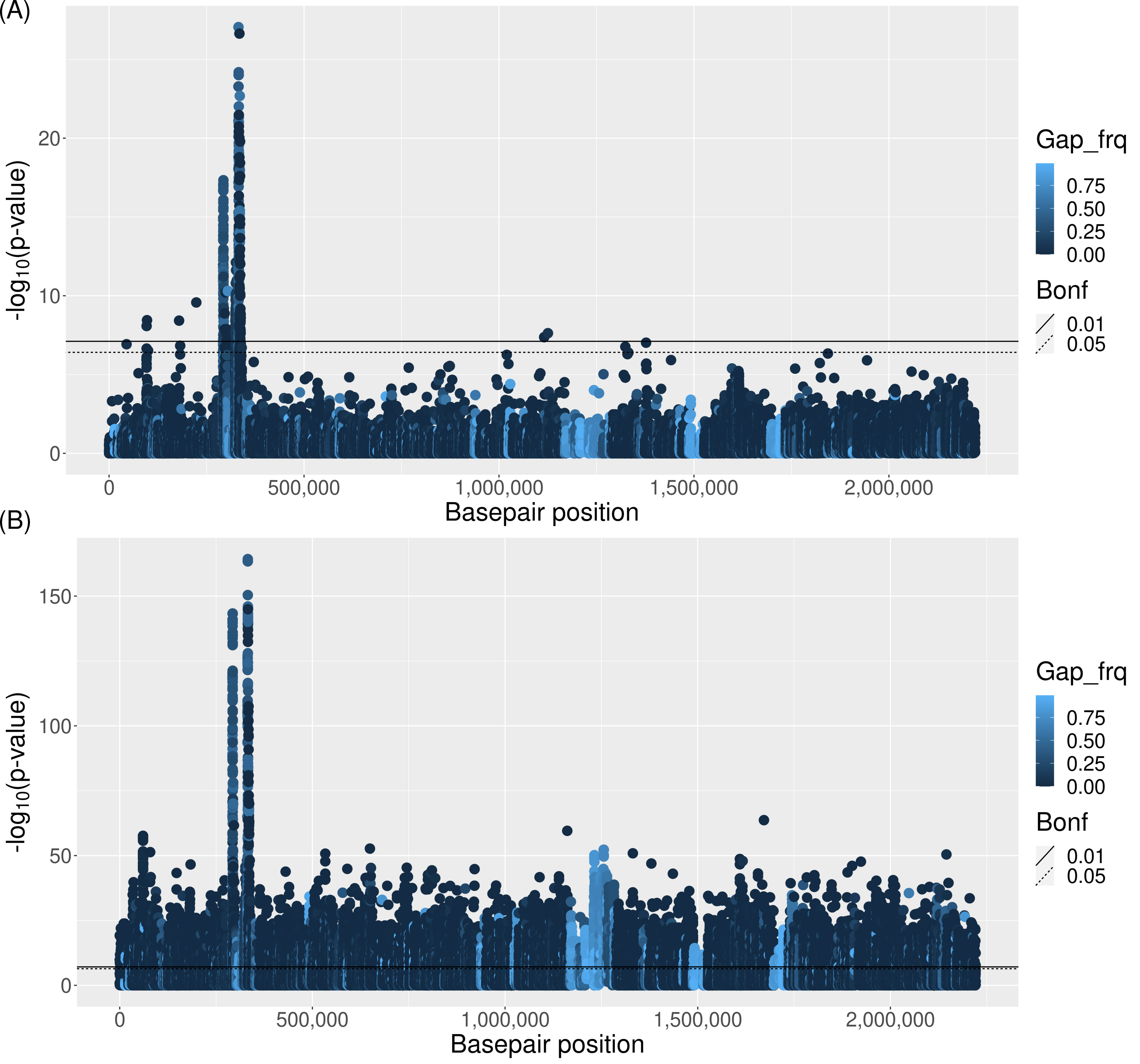

### S12_Fig

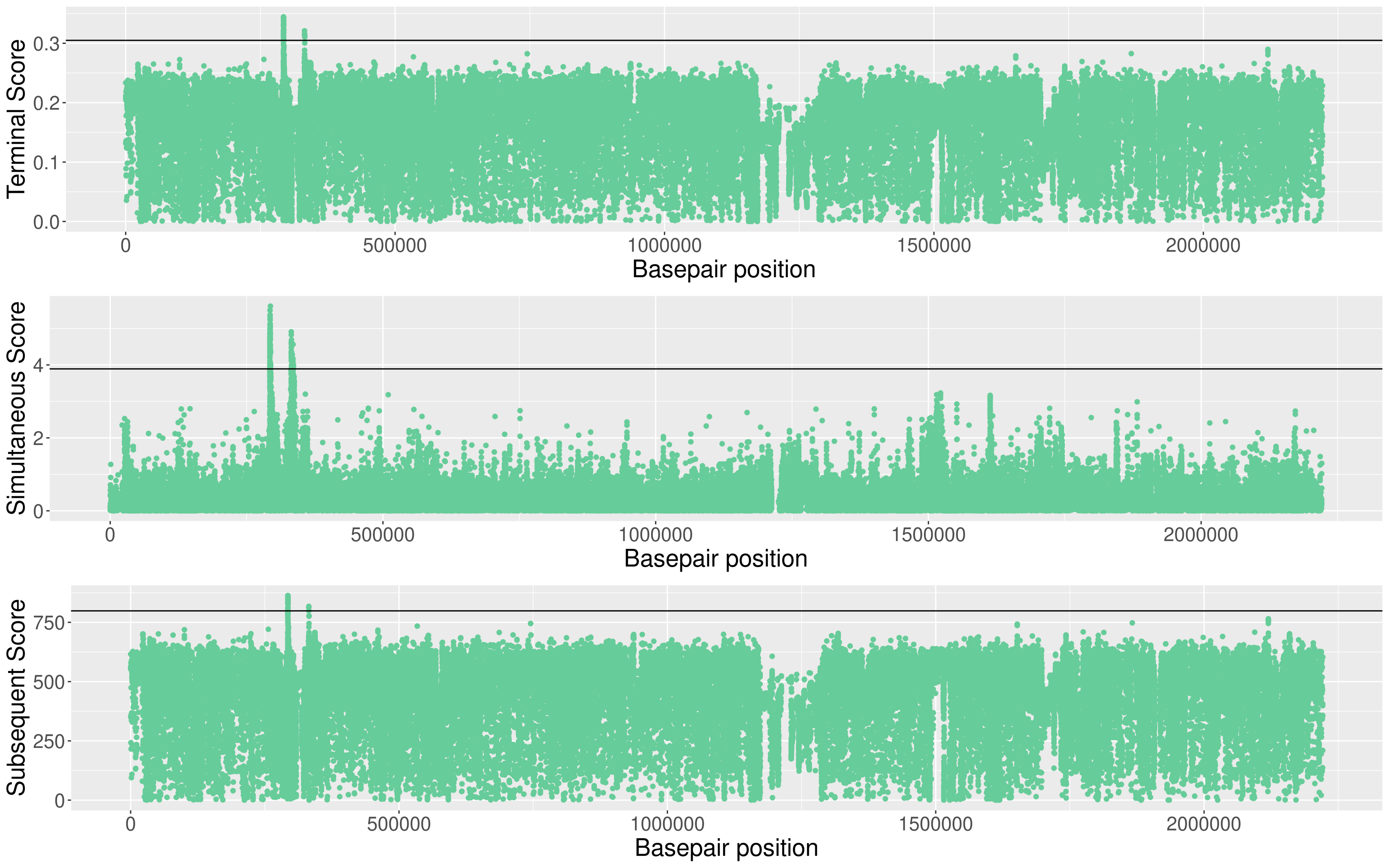

### S13_Fig

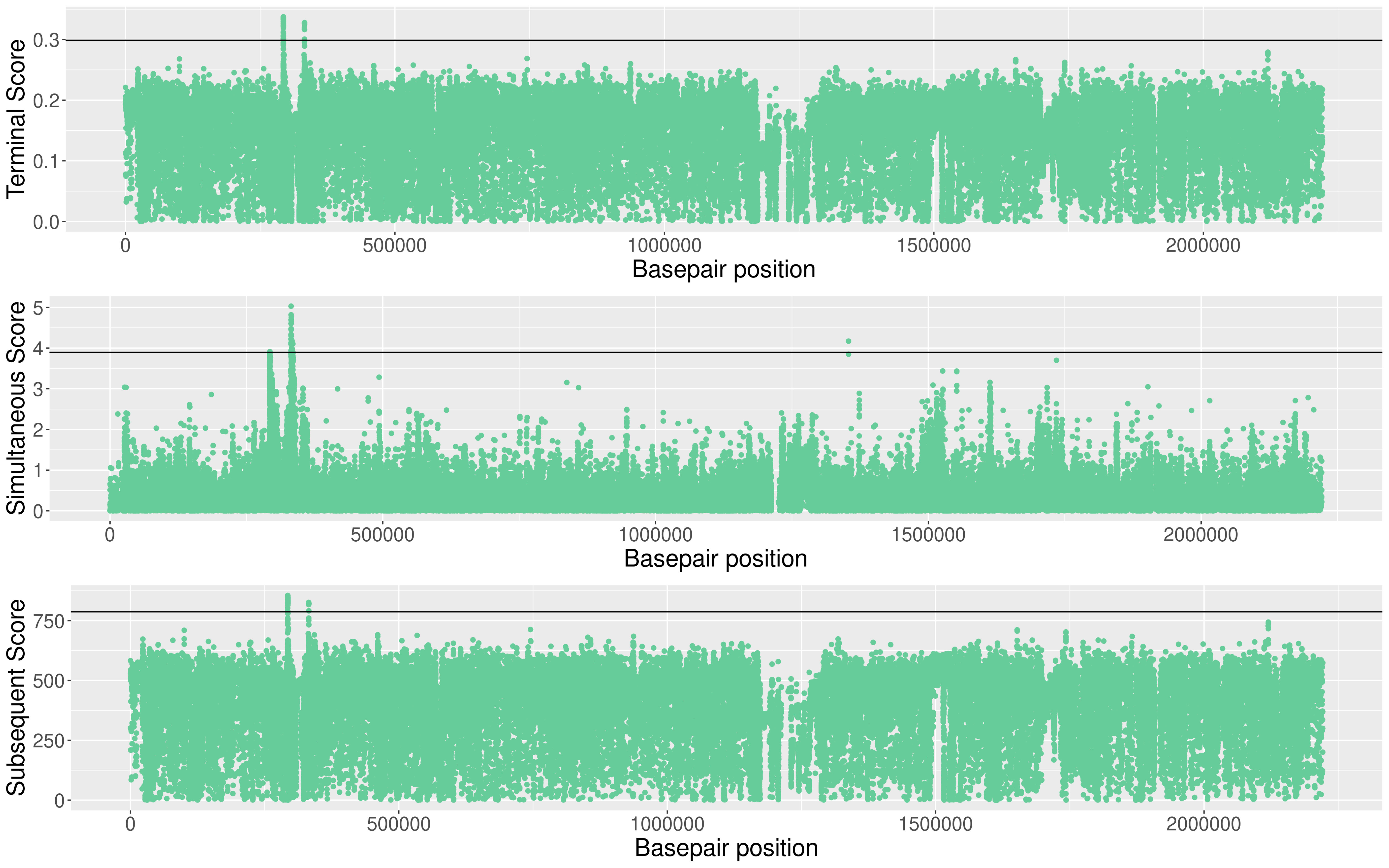

### S14_Fig

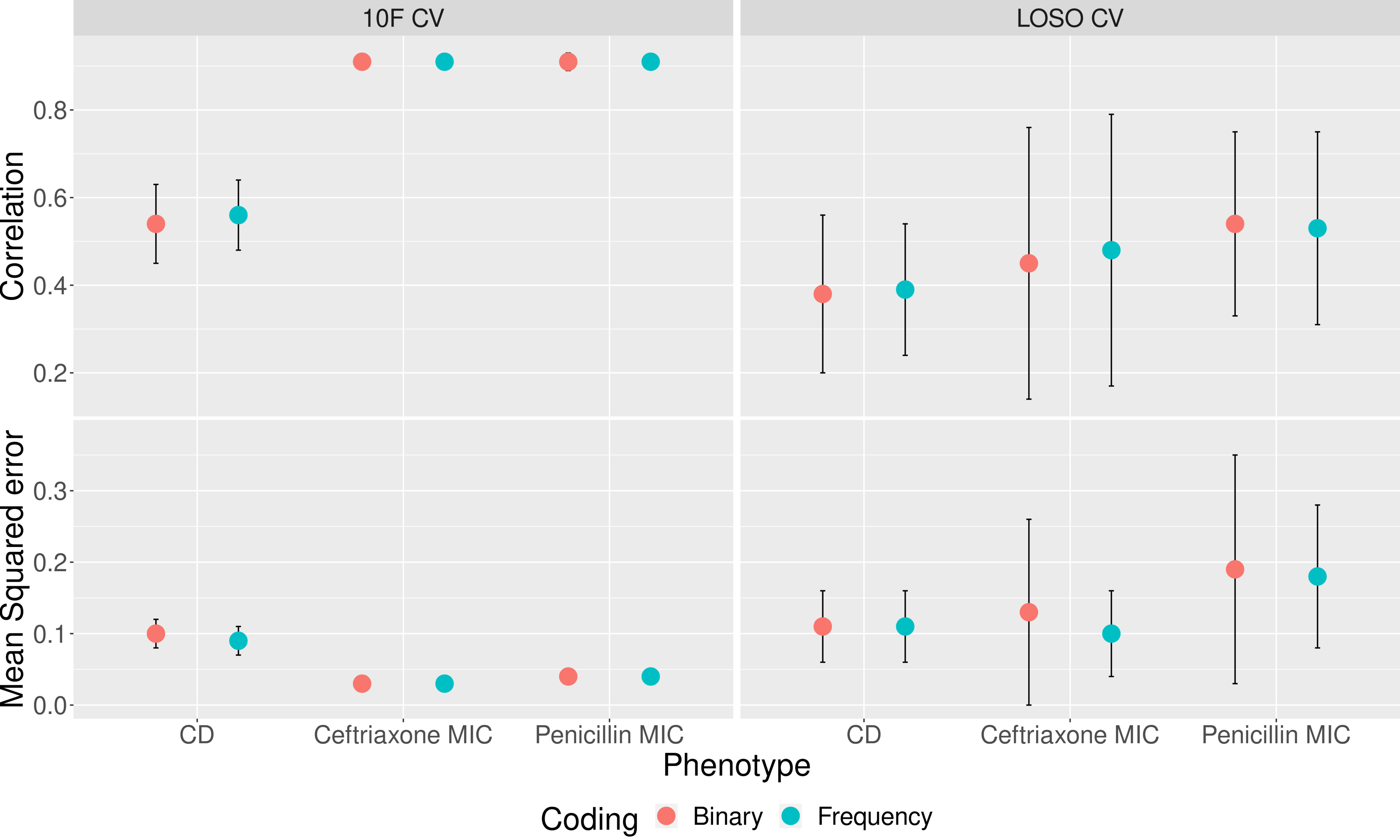
